# Single Nucleus Profiling Highlights the All-Brain Echinoderm Nervous System

**DOI:** 10.1101/2025.03.24.644250

**Authors:** Periklis Paganos, Jack Ullrich-Lüter, Alba Almazán, Danila Voronov, Jil Carl, Anne-C. Zakrzewski, Berit Zemann, Maria Lorenza Rusciano, Tiphaine Sancerni, Maria Schauer, Oğuz Akar, Filomena Caccavale, Maria Cocurullo, Giovanna Benvenuto, Jenifer Carol Croce, Carsten Lüter, Maria Ina Arnone

**Author notes:** these authors contributed equally to this work (Periklis Paganos, Jack Ullrich-Lüter).

## Abstract

Metazoans comprise diverse tissues and cell types, each essential for the survival of the organism. Most of these cell types are established early in embryogenesis and persist into adulthood. However, in indirectly developing sea urchins, the continuity between embryonic and adult stages is thus interrupted by a planktonic larval stage that undergoes complete metamorphosis. In addition, while the gene regulatory networks governing distinct embryonic and larval cell lineages are well studied, the molecular and morphological identities of post-metamorphic sea urchin cell types remain poorly understood. Here, we reconstructed the cell type atlas of post-metamorphic *Paracentrotus lividus* juveniles using single-nucleus transcriptomics, shedding light on the conservation of genetic regulatory mechanisms in post-metamorphic cell types. We identified cell signatures corresponding to eight distinct cell type groups and analyzed at least twenty-nine neuronal cell families, including fifteen unique photoreceptor cell signatures. By combining single cell transcriptomics with spatial gene expression analysis and high-resolution electron microscopy, we identified homologues of vertebrate neuronal genes and photoreceptive opsins expressed throughout the sea urchin body. These findings provide evidence that the echinoderm body plan is not only predominantly head-like, but also exhibits an all-brain organization in an animal previously considered to have a primitive nervous system.

## Introduction

During the last decade, single-cell (scRNA-seq) and single-nuclei (snRNA-seq) RNA sequencing has proven to be an indispensable tool for dissecting the cell type repertoire of a wide range of metazoans. These technologies allow for the comparison of molecular signatures and the investigation of evolutionary origins (1,2). Furthermore, developmental single-cell transcriptomes have enabled the identification of signaling cascades and gene regulatory networks (GRNs) active during embryogenesis, morphogenesis, and organogenesis, revealing previously unknown cell-specific mechanisms (3,4). This revolutionary approach has uncovered dynamic changes in the molecular signatures of cells constituting various cell types, even challenging traditional cell and molecular biology definitions of what constitutes a cell type (5). Recently, significant efforts have been made to identify and characterize cell types by combining omics technologies with morphology-based approaches (6).

Sea urchins are marine deuterostomes featuring a biphasic life cycle with free-swimming bilaterally symmetric planktotrophic or lecithotrophic larvae, followed by a radical metamorphosis leading to animals characterized by a pentaradial adult body plan. Sea urchins have been extensively studied in relation to biological phenomena such as oogenesis, fertilization, embryogenesis, morphogenesis, and organogenesis, at both molecular and GRN levels (7,8). Recently, scRNA-seq studies have focused on understanding the identity and variety of echinoderm cell types in echinoid (9–12) and asteroid (13,14) embryos and larvae. However, these studies were restricted to pre-metamorphic developmental stages, which feature less complex tissue compositions compared to post-metamorphic stages. All post-metamorphic echinoderms share synapomorphies that diversify them from other deuterostomes (15,16). Such examples include their pentaradial symmetrical body plan and their decentralized nervous system organized in a central nerve ring and radial nerve cords along with peripheral nerves. Another one is in regard to their water vascular system that is operating through the podia of echinoderms and is involved in many physiological processes including gas exchange, locomotion and feeding (17).

The GRNs responsible for developing various sea urchin tissues, such as the skeleton, muscles, gut, and nervous system, have been thoroughly investigated in pre-metamorphic stages (18–21) and it has been shown that they produce corresponding (functional) cell types in juvenile and adult animals. Molecular studies on the pre-metamorphic sea urchin nervous system have shown that its composition is far from “primitive,” expressing many genes present in the vertebrate central nervous system (CNS), even during embryogenesis (9, 22–24). However, considerable differences in gene expression profiles and potentially in cell type signatures are expected in post-metamorphic animals, given the vastly different environments that sea urchin larvae and adults inhabit. Different types of photoreceptor cells (PRCs) found in pre- and post-metamorphic sea urchins already exemplify this divergence (23,25–28).

In this study, we investigated the cell type diversity in post-metamorphic juveniles of the sea urchin *Paracentrotus lividus*, with an emphasis on neuronal and PRC types. We used snRNA-seq to generate a cell type atlas, through which we explored cell type identities, examined the conservation of their genetic wiring, and compared their signatures to those found in larval stages. Spatial gene expression and ultrastructural analyses were employed to characterize cell type identity at the (ultra)structural level. Our findings contribute to understanding the evolution and gene regulatory mechanisms controlling development and cell type function in echinoderms and across metazoan phyla.

## Materials & Methods

### Animal husbandry

Adult *P. lividus* individuals were collected from both the Gulf of Naples (Italy) and the Bay of Villefranche-sur-Mer (France) and were housed, respectively, at Stazione Zoologica Anton Dohrn (SZN) and Institut de la Mer de Villefranche (IMEV), in circulating seawater aquaria. At SZN, spawning of gravid animals was induced by vigorous shaking. At IMEV, gametes were collected from dissected gonads. For fertilization, at SZN and IMEV, sperm was diluted 1:1000 and added to a beaker containing unfertilized oocytes in approximately 100 mL of natural filtered sea water (FSW). Fertilization was monitored through microscopic observation and confirmed by the presence of the fertilization envelope. Zygotes were rinsed twice with FSW, to remove the excess of sperm. At SZN, embryos and larvae were reared at a density of 5 embryos/larvae per mL in FSW at 18°C and fed 3 times per week with *Dunaliella tertiolecta.* Twice per week, half of the culture seawater was exchanged with fresh FSW. Upon successful metamorphosis, the juveniles were transferred to new containers, kept at 18°C and water was changed twice per week. Once the mouth was formed juveniles were fed with *Ulva lactuca* algae until they reached the desired stage which was 2 weeks post-metamorphosis (wpm). At IMEV, embryos and larvae were reared at a concentration of, respectively, 70 embryos and 1 larva per mL at 18°C under constant mechanical stirring and fed five times per week with a mixture of *Dunaliella salina* and *Rhodomonas salina*. Three times per week, two-third of the culture sea water was exchanged with fresh FSW. Upon induced metamorphosis using Dibromomethane (29), the juveniles were transferred to new containers and kept at 16°C under constant sea water flow. The sea water was pumped at a depth of 5 m, decanted, and cooled down to 16–18°C. Once the mouth was formed the juveniles were fed with *Tetraselmis suecica* algae until they reached the 2 wpm stage. Juveniles reared at SZN were used for immunohistochemistry, fluorescent *in situ* hybridization, hybridization chain reaction, and electron microscopy experiments. Juveniles reared at IMEV were used for the single nucleus RNA sequencing and hybridization chain reaction experiments.

### Isolation of intact nuclei and single nucleus RNA sequencing (snRNA-seq)

*P. lividus* juvenile nuclei were isolated as previously described (30), with minor modifications. A total of eighteen juveniles was used, issued from two distinct cultures. Three independent snRNA-seq libraries were prepared, each with six 2 wpm juvenile individuals (all from culture 1, all from culture 2 or a 1:1 mix from cultures 1 and 2). All the following steps described here were performed on ice. Juveniles were placed in the lid of a 5 mL Protein LoBind tube (Eppendorf, #0030108302) containing 560 μl of pre-chilled Homogenization Buffer. Homogenization Buffer is a hypotonic buffer composed of 250 mM sucrose, 25 mM KCl, 5 mM MgCl_2_, 10 mM tris buffer (pH 8.0), 1 mM dithiothreitol, 1× protease inhibitor (Sigma-Aldrich, #11873580001), 0.4 U/μl ribonuclease inhibitor (Invitrogen, #AM2682), 0.2 U/μl SUPERase-In ribonuclease inhibitor (Invitrogen, #AM2694), 1 mM spermine, and 0.3% Triton X-100. For each library, the six juveniles were cut in half using a microsurgical knife (Electron Microscopy Sciences, #72047-30), while the tissues from three of them were further gently scraped using a microsurgical blade and fine forceps (Fine Science Tools, #10316-14 and #11251-20, respectively) under a dissecting microscope. Juvenile large fragments were transferred, using a P1000 micropipette, into a clean 1.5 mL Protein LoBind tube. The rest of the supernatant, prior to its transfer into the same tube, was filtered through a 40 μm cell strainer (PluriSelect Mini strainer), to remove debris. Nuclei were extracted through application of mechanical force via pipette aspiration for 10 times using a P1000 micropipette and 15 times using a P200 micropipette. The procedure was performed gently to avoid the formation of bubbles that can lead to nuclei loss or damage. Specimens were left until the large juvenile fragments settled and the supernatant containing the isolated nuclei was transferred into a new clean 1.5 mL Protein LoBind tube. Isolated nuclei were spun down for 10 min at 500 g and 4°C using a 5424 R Eppendorf microcentrifuge. The presence of pellets was visually confirmed (brown colored pellets). All the supernatant was discarded except from 50 μl (surrounding the pellet). 600 μl of Wash Buffer [1x Dulbecco’s phosphate-buffered saline (DPBS), 2% BSA, 0.2 U/μl SUPERase-In ribonuclease inhibitor, 1 mM spermine and 5 mM MgCl_2_] was carefully added (drop by drop) using a P200 micropipette, in order not to disrupt the pellet. Next, the nuclei pellets were resuspended through pipette aspiration for 5 times, ensuring that the resuspended nuclei stay close to the bottom of the tube. Isolated nuclei were once more spun down for 10 min at 500 g and 4°C using a 5424 R Eppendorf microcentrifuge. Most of the supernatant was discarded apart from 25 μl (covering the pellet). Lastly, 25 μl of Resuspension Buffer [3x DPBS, 2% BSA, 0.4 U/μl ribonuclease inhibitor, 0.2 U/μl SUPERase-In ribonuclease inhibitor and 1 mM spermine] were added to the pellet. Pellet was resuspended through gentle pipette aspiration for 20-30 times using a P200 micropipette. Nuclei suspensions were filtered through a 10 μm cell strainer (PluriSelect Mini strainer) to avoid aggregates. The nuclei quality and numbers were estimated by Trypan blue and hemocytometer, respectively. Isolate nuclei (25,000 per library) were loaded on the 10x Genomics Chromium Controller according to the manufacturer’s instructions. cDNA libraries were prepared using the Chromium Single Cell 3’ Reagent Kit (v3.1 Chemistry Dual Index). Libraries were sequenced by the IGFL sequencing platform (Plateforme de Séquençage de l’IGFL) using the Illumina NextSeq 500, and 146 to 171M reads were obtained per sample.

### Mapping of snRNA-seq reads

Prior to mapping, the *P. lividus* genome (31) version 1.0 annotation gene models were extended in the 3’ direction by 5 kilobases (kb), to ensure no overlap with downstream genes as described in (30) and improve mapping rates. Then, the reads were mapped to this genome and modified annotation using the Cell Ranger Software Suite v7.1.0 (32) with --include-introns=true, --no-bam, --nosecondary and --force-cells=N flags, where N is number of cells to force for each sample. This number was determined by mapping the data without forcing a set number of nuclei and visually assessing the Barcode Rank Plot from the web summaries generated by Cell Ranger. This was done in order to ensure that only real nuclei end up in the final analysis. Cell Ranger output matrices, features and barcodes were used for further analysis using the R package Seurat v 4.4.0 (33).

### Analysis of snRNA-seq data

Seurat objects were created by excluding from the analysis genes that are transcribed in less than three cells and cells that have less than a minimum range of 350-500 (depending on the library) and a maximum of 5000 transcribed genes. The generated datasets were normalized, and variable genes were found using the variance stabilizing transfer (VST) method with a maximum of 2000 variable features. The three different objects corresponding to the three independent libraries were integrated through identification of gene anchors (FindIntegrationAnchors) between the different datasets. The integration of the datasets was followed by scaling and principal component analysis (PCA). Jackstraw was used to evaluate PCA significance. A Sharing Nearest Neighbor (SNN) graph was computed with 50 dimensions (resolution 1.0). Uniform Manifold Approximate and Projection (UMAP) was used to perform clustering dimensionality reduction. The final object consisted of 25,000 cells. The FindAllMarkers command was used to identify differentially expressed marker genes. To further confirm *P. lividus* gene IDs, nucleotide BLAST v2.6.0+ (34) searches were performed against *Strongylocentrotus purpuratus* using the -max_target_seqs 1 and - max_hsps 1 options to select a single top hit per *P. lividus* gene sequence and the output setting -outfmt “6 qseqid sseqid” to get both query and subject sequence IDs. Cell type tree reconstruction based on pairwise expression distances between clusters, using all expressed genes, was performed using 10,000 bootstraps as previously described (9). Sub-clustering analysis of the PRCs was performed using the subset function available in Seurat, by using all the clusters corresponding to the nervous system (clusters 1-29) and only the cells that express at least one of the seven opsin genes encoded by the *P. lividus* genome. The subsetted objects were further analyzed by means of calculation of variable genes, scaling, and PCA analysis. Jackstraw was used to determine the number PCAs to be used for clustering. 19 PCAs were selected, and clustering dimensionality reduction was performed. To compare the *S. purpuratus* 3 days post-fertilization (dpf) larva single cell atlas and the *P. lividus* juvenile single nucleus atlas, SAMap v1.02 (35) was used as previously described (36). The average score of ANE, peripheral and neurogenic markers was estimated using the AddModuleScore function incorporated in the Seurat R package.

### Immunohistochemistry (IHC)

Post-metamorphic juveniles were fixed in 4% paraformaldehyde (PFA) in FSW for 20 min at room temperature (RT). Next, specimens were washed once in 100% ice cold methanol (MeOH) for 1 min at RT, followed by multiple rinses with 1x PBS and 0.1% Tween-20 (PBST). Specimens were incubated in a Blocking Solution containing 1 mg/mL Bovine Serum Albumin (BSA) and 4% Sheep Serum (SS) in PBST for 1 hour (h) at RT. Primary antibodies were appropriately diluted in PBST and incubated for 1 h and 30 min at 37°C (or ON at 4°C in the case of Sp-Opsin1). The primary antibodies used were against: Msp130 (gift from Dr. David R. McClay) to label skeletogenic cells (1:20), SYT1 (gift from Dr. Robert Burke) to mark the nervous system (1:20), MHC to label muscles (1:100) (1:100-PRIMM, Italy), and Sp-Opsin1 (1:50) and Sp-Opsin4 (1:50) to mark different photoreceptor types. Following primary antibody incubation, specimens were washed multiple times with PBST and incubated for 1 h with the appropriate secondary antibody (AlexaFluor), diluted 1:1000 in PBST. Sp-Opsin4 IHC paired with Pl-Opsin4 HCR was performed upon the completion of the HCR procedure. The only differences in the IHC protocol in this case are that the fixation step was skipped, and the specimens were kept in the dark throughout the procedure. Specimens were imaged using Zeiss LSM 700 and Leica TCS SP5 confocal microscopes.

### Fluorescent in situ hybridization (FISH)

Fluorescent *in situ* hybridization for *opsin2* was performed as previously described (37,38). In brief, specimens were fixed in 4% PFA in MOPS Buffer (0.1 M MOPS pH 7, 0.5 M NaCl and 0.1% Tween-20 in nuclease-free water) for 1 h at RT, gradually dehydrated and kept at -20°C in 70% ethanol. The signal was developed using the Tyramide Signal Amplification (TSA) technology (Akoya Biosciences) and specimens were imaged on a Zeiss LSM 700 confocal microscope.

### Hybridization Chain Reaction (HCR)

Probes for HCR were generated using the coding sequence of the genes of interest. The *opsin3.2*, *opsin4*, *beta-arrestin*, *vcry*, *otx* and *barh1* probes were synthesized by Molecular Instruments. Probes for, *pax2/5/8*, *delta*, *fzd5/8*, *nkx2.1*, *irxA*, *syt1*, *hdc*, *th*, and *gad* were designed using the custom software developed by (39) and synthesized by Integrated DNA Technologies (IDT). The stock concentration of each probe set was 1 µM. HCRs for *opsin3.2*, *opsin4*, *beta-arrestin*, *vcry*, *otx* and *barh1* were performed at SZN and Museum für Naturkunde (MFN) using a modified protocol from (40), while for *pax2/5/8*, *delta*, *fzd5/8*, *nkx2.1*, *irxA*, *syt1*, *hdc*, *th*, and *gad* HCRs were carried out at IMEV as described in (40,41) with minor modifications. For the HCRs performed at SZN and MFN, the procedure was carried out as follows. 2 wpm juveniles were collected and fixed in 4% PFA in a fixative buffer containing 0.1 M MOPS, 0.5 M NaCl, 2 mM EGTA, 1 mM MgCl_2_ and 1x PBS overnight (ON) at 4°C.The day after, specimens were rinsed three times (5 min each) in fixative buffer at RT and stored in 100% MeOH at -20°C until use. On the day of the HCR experiment, juveniles were rehydrated in decreasing concentrations of MeOH (75%, 50%, 25% in nuclease-free water) and each wash lasted 10 min. Juveniles were washed three times (10 min each) in PBSTr (1x PBS, 0.1% Triton X-100). Specimens were incubated in a decalcification buffer containing 5% EDTA in nuclease-free water for 4 h at RT, and after this step, washed again in PBSTr for three times (5 min each). Juveniles were incubated in permeabilization solution composed of 1% Triton X-100 in 1x PBS at 4°C ON. The day after, the permeabilization buffer was removed, and samples were digested with 4 µg/mL Proteinase K for 6.5 min at 37°C. Samples were washed in PBSTr for two times (5 min each) and post-fixed in 4% PFA for 1 h at RT. Samples were washed again for five times (5 min each) with PBST and afterwards incubated in the probe hybridization buffer (Molecular Instruments) at 37°C for 2 h. Within this time interval, probe hybridization buffer was exchanged once. Probe solution was prepared by diluting each probe in the probe hybridization buffer to a final concentration of 0.04 μM. Specimens were incubated with the probe solution ON at 37°C. Following the probe hybridization, specimens were washed three times (5 min each) and two times (30 min each) with pre-heated probe wash buffer (Molecular Instruments). Specimens were washed several times with 5x SSCTr (5x saline-sodium citrate (SSC), 0.1% Triton X-100). A pre-amplification step was performed by incubating the sample for 30 min at RT in an equilibrated to RT amplification buffer (Molecular Instruments). In the meantime, 2 μl of each hairpin from Molecular Instruments (of 3 µM stock) were placed individually in tubes. The hairpins h1 and h2 were heated up separately by placing the tubes at 95°C for 90 seconds. After this step, the hairpins were stored on ice, in the dark, for 30 min. The hairpins were then added to the amplification buffer at a final concentration of 0.06 µM. The specimens were incubated in the hairpin solution at RT, in the dark, for at least 20 h. The following day, the samples were washed several times in 1x SSCTr. The juveniles were transferred to a solution containing 1 µg/mL DAPI in 1x SSCTr, then transferred to 50% glycerol (in 1x PBS, pH 7.4) and mounted for imaging. Images were acquired using Zeiss LSM 700 confocal and Leica TCS SP5 microscopes.

At IMEV, HCRs were carried out as follows. 2 wpm juveniles were fixed ON at 4°C in 4% PFA, 10 mM EPPS diluted in FSW. Upon fixation, juveniles were washed once for 5 min at RT in 1x PBST and once for 5 min at RT in cold (-20°C) MeOH, before being stored at -20°C in MeOH. Fixed juveniles were rehydrated through progressive washes in 75%, 50%, 25% MeOH in 1x PBST (5 min each) at RT. Juveniles were then washed twice (10 min each) at RT in 100% 1x PBST, followed by three washes (10 min each) at RT in RNase free water. For tissue clearing purposes, juveniles were incubated ON at 4°C in 50% tetrahydrofuran (THF) (#186562, Sigma-Aldrich, Saint-Quentin-Fallavier, France) in RNase free water (42). The THF was thereafter discarded by three washes (30 min each) at RT in RNase free water and two washes (5 min each) at RT in 1x PBST. Permeabilization of the juveniles was conducted by incubation in detergent solution, containing 50 mM TrisHCL (pH 7.5), 1 mM EDTA, 150 mM NaCl, 1% SDS, 0.5% Tween-20, for 1 h at RT. Juveniles were washed four times (5 min each) at RT in 1x PBST, before they were transferred into 96-well plates and prehybridized for 3 h at 37°C in 50 µL of hybridization buffer containing 30% formamide, 5x SSC, 9 mM citric acid (pH 6.0), 50 µg/mL heparin, 1× Denhardt’s solution, 10% dextran sulfate, 0.1% Tween-20. During the pre-hybridization step, each probe pair set was diluted in 50 µL of hybridization buffer, to a final concentration between 0.008 nM to 0.04 µM depending on the targeted gene. Probe pair dilutions were then added to the specimens and placed in a humid chamber ON at 37°C to allow probe pairs hybridization. Probe pairs were washed out by four washes (5 min each) at 37°C, and then two washes (30 min each) at 37°C, all using pre-heated probe wash buffer (30% formamide, 5× SSC, 9 mM citric acid (pH 6.0), 50 µg/mL heparin, 0.1% Tween-20). Next, two additional washes were performed (5 min each) at RT in 5x SSCT (5x SSC, 0.1% Tween-20), followed by two washes (5 min each) at RT using 1x SSCT. Juveniles were subsequently incubated in 50 µL of amplification buffer (5x SSC, 10% dextran sulfate, 0.1% Tween-20) for 30 min at RT. During the pre-amplification step, h1 and h2 hairpins (Molecular Instruments) were prepared as described above for HCRs in SZN and MFN, except that only 1 µL of each hairpin (3 µM stock) was used. Upon cooling down, h1 and h2 hairpins were transferred together in 50 µL of amplification buffer. Diluted hairpins were added onto the specimens and incubated ON in the dark at RT. Hairpin removal was carried out by several washes, all performed in the dark at RT. These washes included two washes (5 min each) in 1x SSCT, one wash (30 min) in 1x SSCT containing 1:1000 Hoechst 33342 for nuclei staining, two washes (30 min each) in 1x SSCT, and four washes (5 min each) in 1x PBST. Juveniles were mounted in an EZ clear mounting medium prepared as described by (42). Images were acquired on a Leica SP8 confocal microscope.

### Transmission electron microscopy (TEM)

Post-metamorphic juveniles were fixed in 1.25% glutaraldehyde in ASW + 1% osmium at 18°C for 1 hour at room temperature (RT). Samples were subsequently washed in PBS (0.05 M phosphate/0.3 M NaCl), pH 7.4, for 10, 30, and 60 minutes, then cooled to 4°C and washed again after 4 hours and overnight. The samples were post-fixed in 1% osmium/1.5% ferrocyanide in 0.2 M cacodylate buffer for 60 minutes at RT. Following two 15-minute washes in distilled water, samples were incubated in 0.3% tetracaine hydrochloride (TCH) in 0.2 M cacodylate buffer (freshly prepared from a 1% TCH stock solution) for 15 minutes. Samples were then washed twice in distilled water for 15 minutes each, followed by incubation in 1% osmium in cacodylate buffer for 60 minutes. After two additional 15-minute washes in distilled water, samples were dehydrated in a graded ethanol series (30%, 50%, 70%, 80%, 90%, 95%) for 10 minutes each, followed by three 15-minute washes in 100% ethanol. After post-fixation, specimens were dehydrated in a graded alcohol series and embedded in plastic resin (Araldite M, Fluka; art. no.: 10951) for 48 hours at 60°C. Serial sections were cut using a diamond knife and a Reichert ultramicrotome S and stained with uranyl acetate and lead citrate (Phoenix Ultrastain, Staining Technologies). Examination was carried out using a ZEISS EM 900 transmission electron microscope (TEM).

### Serial block face scanning electron microscopy (SBF-SEM)

Two-week post-metamorphic (*wpm*) *Paracentrotus lividus* juveniles were collected and fixed on ice for 1 hour in 1.25% glutaraldehyde and 1% osmium in artificial seawater (ASW). After fixation, samples were kept on ice and washed with cold PBS (0.05 M phosphate buffer/0.3 M NaCl, pH 7.4) for 10 minutes, 30 minutes, 4 hours, and overnight (ON) at 4°C. All post-fixation steps were performed at room temperature (RT). Samples were incubated in 0.2 M cacodylate buffer with 1% osmium and 1.5% ferrocyanide for 1 hour. The specimens were then rinsed twice in PBS (15 minutes each) and incubated for 15 minutes in 0.3% tetracaine hydrochloride (TCH) in 0.2 M cacodylate buffer, followed by another two PBS washes. Subsequent incubation in 1% osmium in 0.2 M cacodylate buffer for 1 hour was followed by two final PBS washes before samples were transferred to distilled water. For decalcification, samples were incubated in a 1:1 mixture of 2% ascorbic acid and 0.3 M NaCl at RT until the skeleton was completely dissolved. The samples were then washed with distilled water, dehydrated in a graded alcohol series, and embedded in plastic resin (Araldite M, Fluka; art. no.: 10951) for 48 hours at 60°C. Acquisition was performed at the Electron Microscopy Core Facility (MCFE) at the European Molecular Biology Laboratory (EMBL), Heidelberg, Germany. Targeting of the region of interest prior to serial block-face scanning electron microscopy (SBF-SEM) acquisition was accomplished using X-ray imaging (Bruker Skyscan 1272). Target region trimming was performed using a Leica UC7 ultramicrotome. Imaging was carried out using a Zeiss GeminiSEM 450 equipped with a Gatan 3View system and an OnPoint backscattered electron detector, using an acceleration voltage of 1.5 kV and a current of 350 pA. The 3View microtome was set to remove 40 nm from the block surface at each cutting iteration. After each cut, a low-resolution overview image of the entire block face was acquired, along with a high-resolution tiled image of the region of interest. The latter was acquired with a 10 nm pixel size and a dwell time of 1.6 µs. To cover the entire region of interest, 6 to 10 tiles (4096 × 3072 pixels) were acquired per section, with an overlap of 350 pixels. The subsequent alignment of tiles resulted in a dataset of serial images of the region of interest with a voxel size of 10 nm × 10 nm × 40 nm. Segmentation and analysis were performed using the software package Amira (Thermo Fisher Scientific), Versions 2021.2 & 2022.2. Nuclei were segmented semi-automatically using various tools in Amira, such as “Thresholding” and “Magic Wand.” Plasma membranes and other cell organelles were segmented manually. The visual display of all segmented reconstructions was achieved using “Surface Rendering” within the software package.

## Results

### Cell type identity and diversity in *Paracentrotus lividus* juveniles

Sea urchin embryos and larvae have been extensively used as research subjects to decipher key developmental processes and to identify GRNs that drive the formation of distinct cell types, many of which are either evolutionary conserved or provide missing links essential for understanding the evolution of complex organs. Nowadays, performing single cell transcriptomics on echinoderm pre-metamorphic stages is a trivial process and single cell RNA sequencing (scRNA-seq) and single nuclei RNA sequencing (snRNA-seq) atlases have been reconstructed for several echinoderm species. However, very little is known regarding the cell types of the post-metamorphic sea urchin juvenile, and one reason for this is the difficulty to extract intact single cells primarily due to the presence of the extensive biomineralized endoskeleton.

To bridge this gap, we devised an efficient single nucleus extraction protocol (detailed protocol in materials and methods) and performed snRNA-seq on 2 wpm *P. lividus* juveniles (Fig. 1A-C). The choice of that developmental stage was based on the fact that 2 weeks post metamorphosis allows enough time for ensuring the absence of any larval tissue remnants and the opening of the mouth and functionality of the adult digestive tract, thereby increasing the probability of highly diversified cell types to be present. Altogether, we constructed three snRNA-seq libraries, each containing transcripts corresponding to nuclei originating from six juveniles per library, that emerged from two genetically diverse animal cultures. Once the libraries were sequenced and mapped, computational analysis was carried out in RStudio using the Seurat pipeline. To merge the three different snRNA-seq libraries, anchor-based integration was performed and resulted in the generation of the 2 wpm *P. lividus* juvenile cell type atlas displayed in Figure 1. Our juvenile integrated atlas is composed of 25,000 nuclei (5,000 nuclei from library 1, 10,000 from library 2 and 10,000 from library 3) distributed to forty-eight clusters, with cells from all three libraries contributing to the formation of the integrated cluster. Each one of the reconstructed clusters corresponds to either distinct cell types, or to related cell type groups (Fig. 1, Fig. S1A-B).

**Fig. 1.**
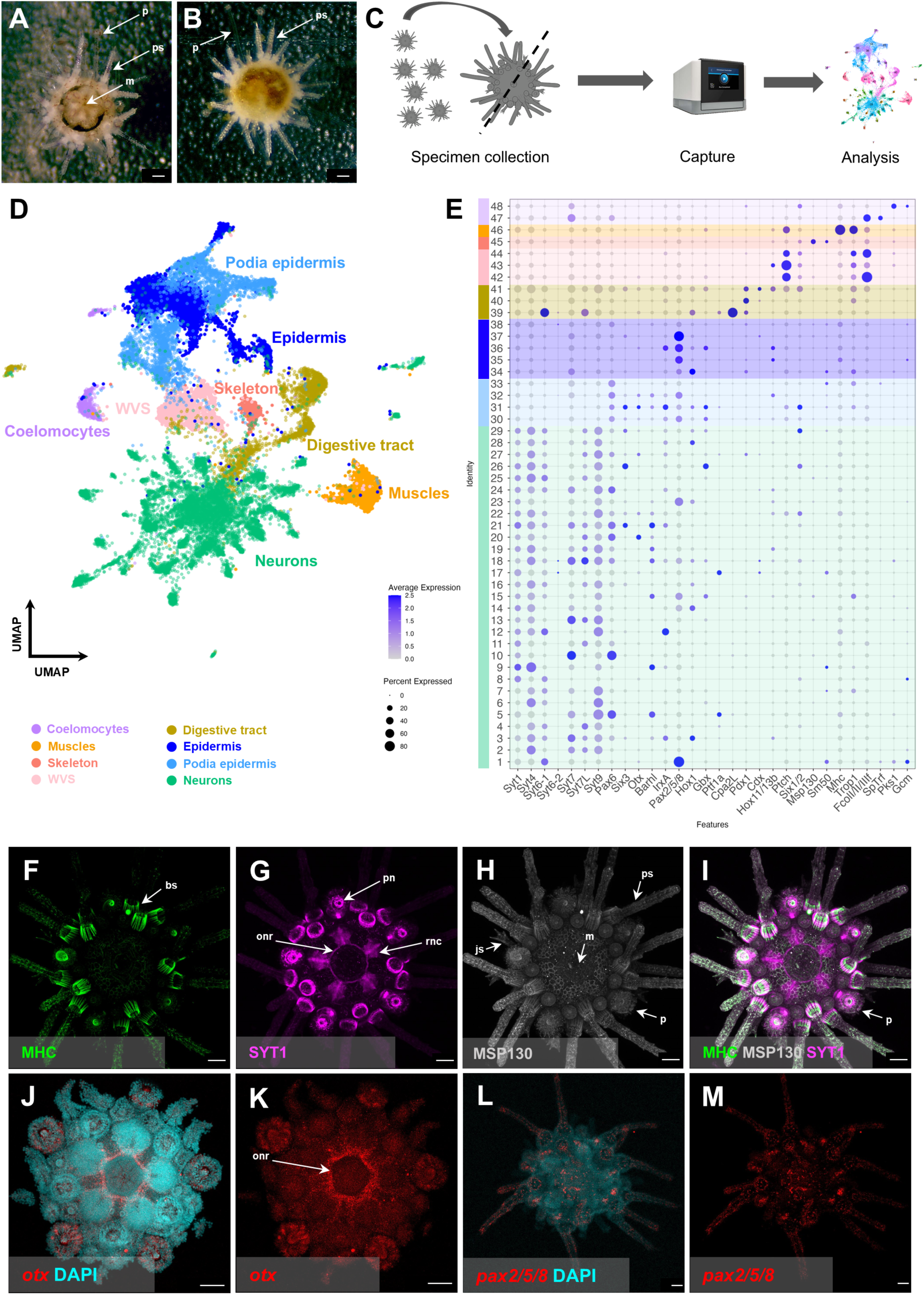
Cell type diversity of the *P. lividus* 2 wpm juvenile. (**A**) 2 wpm *P. lividus* juvenile seen from the oral side. (**B**) 2 wpm *P. lividus* juvenile seen from the aboral side. (**C**) Schematic representation of our snRNA-seq pipeline: from intact juveniles to single nuclei, sequencing and computational analysis. (**D**) Integrated UMAP of the three snRNA-seq libraries. Cells are color-coded in respect to the cell type groups recognized. (**E**) Dotplot showing the average expression of genes used to assign cluster identities. Color-code is the same as in panel D. (**F**) IHC for myosin heavy chain (MHC), labeling muscles. (**G**) IHC for synaptotagminB (SYT1), labelling neurons. (**H**) IHC for MSP130, labeling skeleton. (**I**) Overlay of MHC, SYT1 and MSP130. (**J-K**) HCR for *otx* with (**J**) and without (**K**) nuclei labelling (DAPI). (**L-M**) HCR for *pax2/5/8* with (**L**) and without (**M**) nuclei labelling (DAPI). (**F-M**) Juveniles are in oral view. bs, base of the spine; m, mouth; onr, oral nerve ring; p, podia; pn, podia neurons; js, juvenile spine; ps, primary spine; rnc, radial nerve cord. Scale bar is 50 μm.

To discover the identity of the generated clusters we calculated the marker genes of each of the clusters and took advantage of i) the extensive echinoderm literature providing us with a great number of gene markers known to label distinct embryonic and larval sea urchin cell types, ii) a (limited) number of known gene markers marking sea urchin juvenile cell types, and iii) a gene marker toolkit originating from a recent publication describing tissue-specific gene expression patterns in post-metamorphic juveniles of the sea star *Patiria miniata* (40). Plotting for the average expression of: neural genes (*syt1, syt4, syt6-1, syt6-2, syt7, syt7L* and *syt9*) (9,13,22); genes labeling the podia epithelium (*pax6, six3, otx, barhl and irxA*) (37,40); epidermal genes (*pax2/5/8, hox1, gbx*) (40); pancreatic acinar-like genes (*ptf1a* and *cpa2L)* (43); posterior gut genes (*pdx1, cdx* and *hox11/13b*) (44); water vascular system (WVS) enriched genes (*ptch* and *six1/2*) (40); skeletal genes (*msp130* and *sm50*) (45); muscle genes (*mhc* and *trop1*) (46,47), and immune system genes (*fcolI/II/IIIf, spTrf, pks1* and *gcm*) (9,48) we were able to group the clusters into eight distinct cell type groups corresponding to neurons, epidermis, podia epidermis, digestive tract, muscles, coelomocytes, water vascular system (WVS) and skeleton (Fig. 1D-E).

In order to perform unbiased reconstruction of the transcriptional relationships of the different clusters, we further performed a cluster-tree analysis (Fig. S2). This analysis showed that most of the clusters corresponding to the nervous system form a well-defined group that contains the coelomocyte cluster (48) as an outlier. Moreover, the skeletal, WVS and muscle clusters along with the coelomocyte cluster 47 are grouped together, reflecting potentially common mesodermal origins. Lastly, the three digestive tract clusters form a well-supported group, as well as clusters corresponding to general and podia epidermis, except for the podia epidermis cluster 31.

To confirm the snRNA-seq analysis predictions and our cluster annotation, we next performed immunohistochemistry (IHC) and hybridization chain reaction (HCR) to assess the spatial profiles of several of the genes included in our analysis (Fig. 1F-M). For instance, we carried out IHC on the sea urchin juveniles using sea urchin antibodies against Syt1 (formerly named SynB for synaptotagminB), myosin heavy chain (MHC) and Msp130 known to label, respectively, the entire nervous system (including the oral nerve ring (ONR), the radial nerve cords (RNCs) and the podia neurons), musculature and skeleton of the sea urchin juvenile (Fig. 1F-I). We further conducted HCR for *otx*, predicted by our analysis to be expressed in clusters corresponding to the nervous system, and *pax2/5/8*, predicted to be expressed in the nervous system, the epidermis and the podia of the juvenile. As expected, the IHC against Syt1 showed immunoreactivity in a massive portion of the juvenile body, in positions formerly identified as the ONR, the RNCs, and the nervous systems associated with the WVS and the spines (Fig. 1G, I) (37,49). Likewise, the IHC for MHC highlighted the muscles in *P. lividus* 2 wpm juveniles both at the level of the spines (Fig. 1F, I) and within the body cavity, as previously reported (45,49), and the same applies for MSP130 outlining the skeleton (Fig. 1H, I) (45). For *otx* and *pax2/5/8*, we detected their transcripts, for both, in presumptive nervous system cells (Fig. 1K-M), with *otx* being enriched in the ONR and the podia nervous system, while *pax2/5/8* being predominantly expressed in neurons related to the spines and podia.

Noteworthy, looking at the distribution of cells per cluster we detected that the most cell-enriched clusters correspond to epidermal and neuronal cells, while the nervous system itself is represented by twenty-nine clusters out of the forty-eight total ones (Fig. S1C).

Once the identity of the cell types was established, we set out to explore how similar the molecular signature of some of those cell types is when compared to the larval ones. To do so, we plotted for the average expression of genes that are either essential components of known sea urchin embryonic or larval GRNs or patterning genes giving rise to diverse cell types (Fig. 2).

**Fig. 2.**
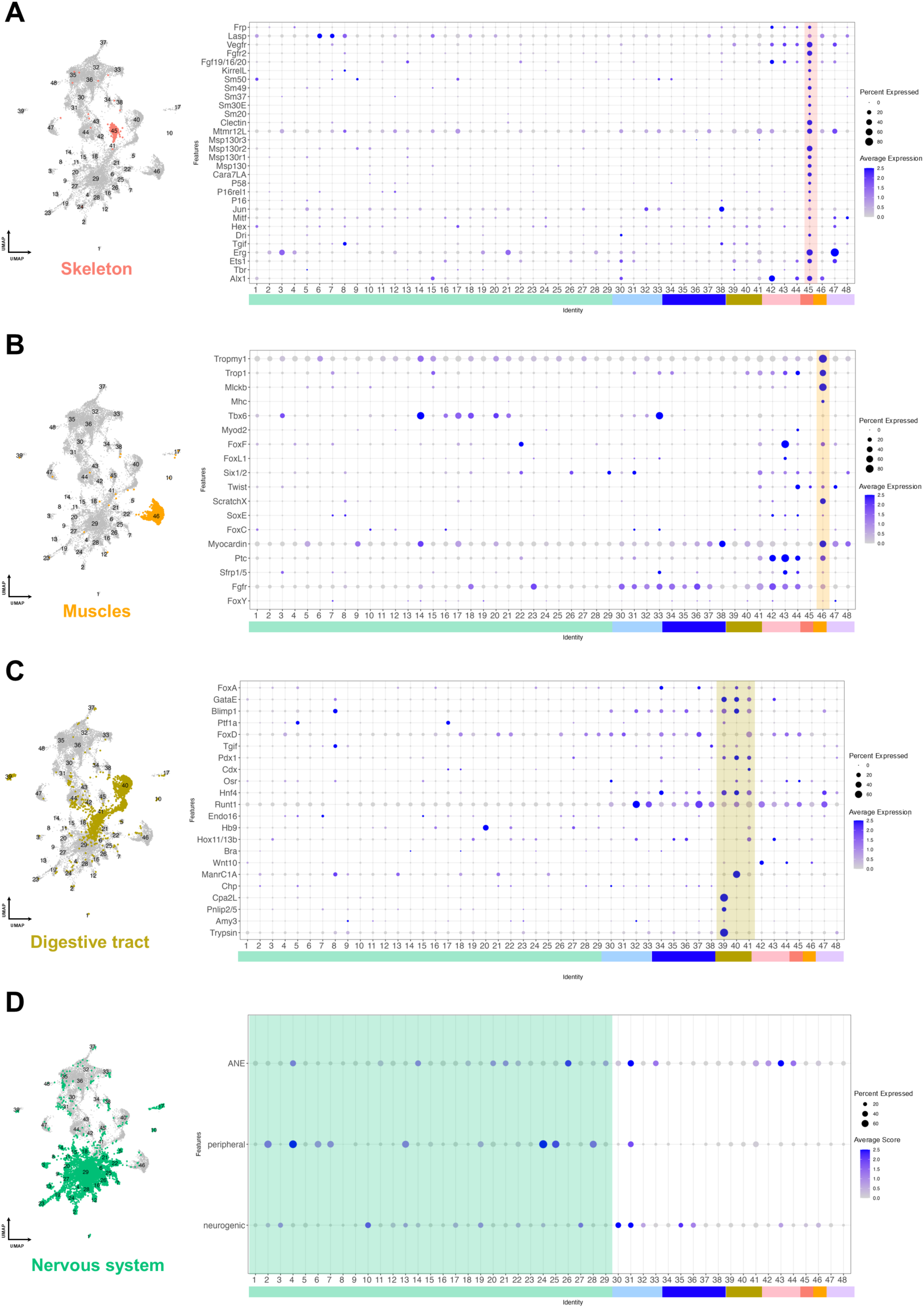
Regulatory signatures of the *P. lividus* 2 wpm juvenile cell types. (**A**) UMAP highlighting the skeletal cluster in the *P. lividus* snRNA-seq data and dotplot showing the average expression, in the *P. lividus* snRNA-seq atlas, of genes constituting the *S. purpuratus* larval skeletogenic GRN. (**B**) UMAP highlighting the muscle cluster in the *P. lividus* juvenile snRNA-seq data and dotplot showing the average expression, in the *P. lividus* juvenile snRNA-seq atlas, of gene markers part of the *S. purpuratus* larval myogenesis GRN. (**C**) UMAP highlighting the digestive tract clusters in the *P. lividus* juvenile snRNA-seq data and dotplot showing the average expression, in the *P. lividus* snRNA-seq atlas, of genes patterning different domains of the *S. purpuratus* larval gut. (**D**) UMAP highlighting the neuronal clusters in the *P. lividus* juvenile snRNA-seq data and dotplot showing the average score, in the *P. lividus* snRNA-seq atlas, of genes expressed in the sea urchin embryonic ANE, embryonic and larval peripheral neurons or necessary for embryonic neurogenesis. Color code is the same as in Fig.1 E.

First, we interrogated our snRNA-seq data for the expression of the genes constituting the skeletogenic GRN, one of the best characterized echinoderm GRNs to date (Fig. 2A). It has previously been postulated that elements of the skeletogenic GRN are reutilized in the post-metamorphic adult program (50). Plotting for the average expression of the skeletogenic transcription factors and the differentiation gene battery, we found that, with the exception of the three transcription factors *tbr, tgif, hex,* and the terminal differentiation spicule matrix/C-lectin domain family member *msp130r3*, all skeletal embryonic and larval genes are operating in juvenile skeletogenic cells, indicating that the juvenile skeletogenic program is highly similar to the embryonic and larval one (Fig. 2A).

Second, investigating the expression profile of genes involved in the GRN controlling the specification and differentiation of sea urchin embryonic myoblasts (20), we found that core regulatory genes, including *foxY, fgfR, sfrp1/5, ptc, myocardin, foxC, soxE, scratchX, twist, foxL1, foxF* and *myod2* are expressed in the cluster corresponding to the juvenile muscles, as well as the differentiation genes corresponding to *mhc, mlckb*, tropomyosin and troponin1 (Fig. 2B). Interestingly, we also found that many of these genes further acquired a potential novel role, as we found them to be expressed in cell types outside of the muscle lineage, such as in the nervous system and the WVS (Fig. 2B).

In the case of the GRNs controlling the diversification of different larval gut cell types and domains, we found their components distributed across the three digestive tract juvenile clusters (39 to 41) (Fig. 2C). In detail, cluster 39 corresponds to the pancreatic acinar-like cells, as the typical larval gene markers of that cell type (43), including the transcription factors *ptf1a, gataE, hnf4*, and *blimp1*, and the terminal differentiation genes *cpa2L, pnlip2/5, amy3*, and *trypsin*, were found co-expressed there. Interestingly, we also report the expression of the pancreatic transcription factor *pdx1* in the acinar-like cells, a domain of expression that has not been reported so far in the sea urchin larval gut. Cluster 40 matched the transcriptomic profile of the larval midgut domain expressing genes such as *foxA, gataE, blimp1, ptf1a, tgif, osr, cdx, pdx1, hnf4, runt1* and *manrC1A* (9,44). Lastly, cluster 41 resembles the embryonic and larval posterior gut, including the pyloric sphincter, the intestine and the anus, by expressing genes such as *foxA, gataE, blimp1, foxD, pdx1, cdx, osr, hnf4, hb9* and *hox11/13b* (9,44).

Finally, we explored the expression profile of transcription factors and signaling molecules that are essential for embryonic and larval sea urchin neurogenesis (9,21,51). Moreover, we scored for the average expression of genes necessary for the specification of the anterior neuroectoderm (ANE) and for the differentiation of the serotonergic neurons that arise from that neurogenic domains as well as for genes expressed in the larval peripheral nervous system constituted by the ciliary band and esophageal neurons. Strikingly, we found most of the genes tested to be expressed by cells belonging to neuronal clusters, suggesting that juvenile neurogenesis is reusing the transcription factor repertoire operating during embryonic and larval development (Fig. 2D, Fig. S3I). Outside the nervous system we find all of the three gene groups present in different clusters corresponding to podia epidermis, while the groups composed of genes involved in the embryonic specification of ANE and neurons were also found in the WVS and epidermis cell type groups, respectively. To our surprise we found very little overlap between the three different groups scored, suggesting a diversification of neuronal fate, similar to the embryonic and larval one. Interestingly the ANE associated genes were found to be differentially expressed in 9 out of the total 29 neuronal clusters, with the highest score found in cluster 26, and the most widespread distribution in cluster 29, which according to our snRNA-seq atlas contains the highest number of cells. Of the ANE genes tested, *six3, fezf, rx, frz5/8* and *nkx2.1,* all members of an evolutionary conserved neurogenic anterior GRN in deuterostomes (52), were differentially expressed in the juvenile nervous system, while absent from the non-neuronal ANE positive clusters. To test our predicted expression patterns, we performed HCR for the neurogenic marker *delta,* the ANE markers *frz5/8* and *nkx2.1* and the transcription factor *irxA* predicted by our data to be expressed in juvenile neurons (Fig. 1E). Our HCRs showed that *delta, frz5/8* and *nkx2.1* are expressed indeed in nuclei within the ONR and RNC regions, while *irxA* was more confined within the RNC portion of the nervous system (Fig. S3).

Following our gene candidate-based analysis, we also used the SAMap pipeline to compare our *P. lividus* snRNA-seq atlas to the already available scRNA-seq atlas of the *S. purpuratus* 3 dpf larva (Fig. 3A). This analysis allowed us to assess cell type conservation between the two developmental stages of the two distinct species, in a holistic and unbiased way. We observed that juvenile muscle, skeletal, and digestive tract clusters (respectively, clusters 46, 45, and 39-40) align with the respective ones of the *S. purpuratus* larva, further supporting our previous analysis (Fig. 3A-C). Interestingly, the larval esophagus, intestine and anus do not align with any of the juvenile ones. Moreover, we found that juvenile coelomocyte clusters 47 and 48 align with the larval blastocoelar and immune cells, respectively. The larval blastocoelar cells also aligned with two juvenile WVS clusters (42 and 44). We also found a population of juvenile neurons (cluster 10) aligning with the larval cluster corresponding to the aboral ectoderm. The aboral ectoderm larval cluster also aligned with three juvenile general epidermis clusters (35–37) and two juvenile podia epidermis clusters (32 and 33). The three general epidermis clusters (35–37) also aligned with the larval ciliary band and apical plate clusters, although with lower alignment scores. Likewise, the two juvenile podia epidermis clusters (32 and 33) aligned with the larval upper and lower oral ectoderm clusters. Interestingly, although we previously found expression of the larval neurogenic patterning genes in the juvenile nervous system (Fig. 2D), the SAMap analysis failed to find any cluster alignment between the juvenile and larval neuronal clusters, suggesting that while the same genes are used the resulting neuronal cell types are vastly different. Noteworthy, our analysis also failed to detect similarities between any of the juvenile clusters and the larval coelomic pouches, a domain where a germ cell-like molecular signature has been previously reported. (Fig. 3). However, genes typical of germ cells and stem cells (e.g., *vasa, piwi, nanos, boule, tudor, msy*) were found expressed throughout the snRNA-seq atlas in various clusters (Fig. S4). The lack of co-expression of these genes in a specific and defined cluster suggest the absence, at 2 wpm of a committed germline lineage, and propose that these genes, at this stage, either mark stem cell-like populations or are incorporated in somatic regulatory programs.

**Fig. 3.**
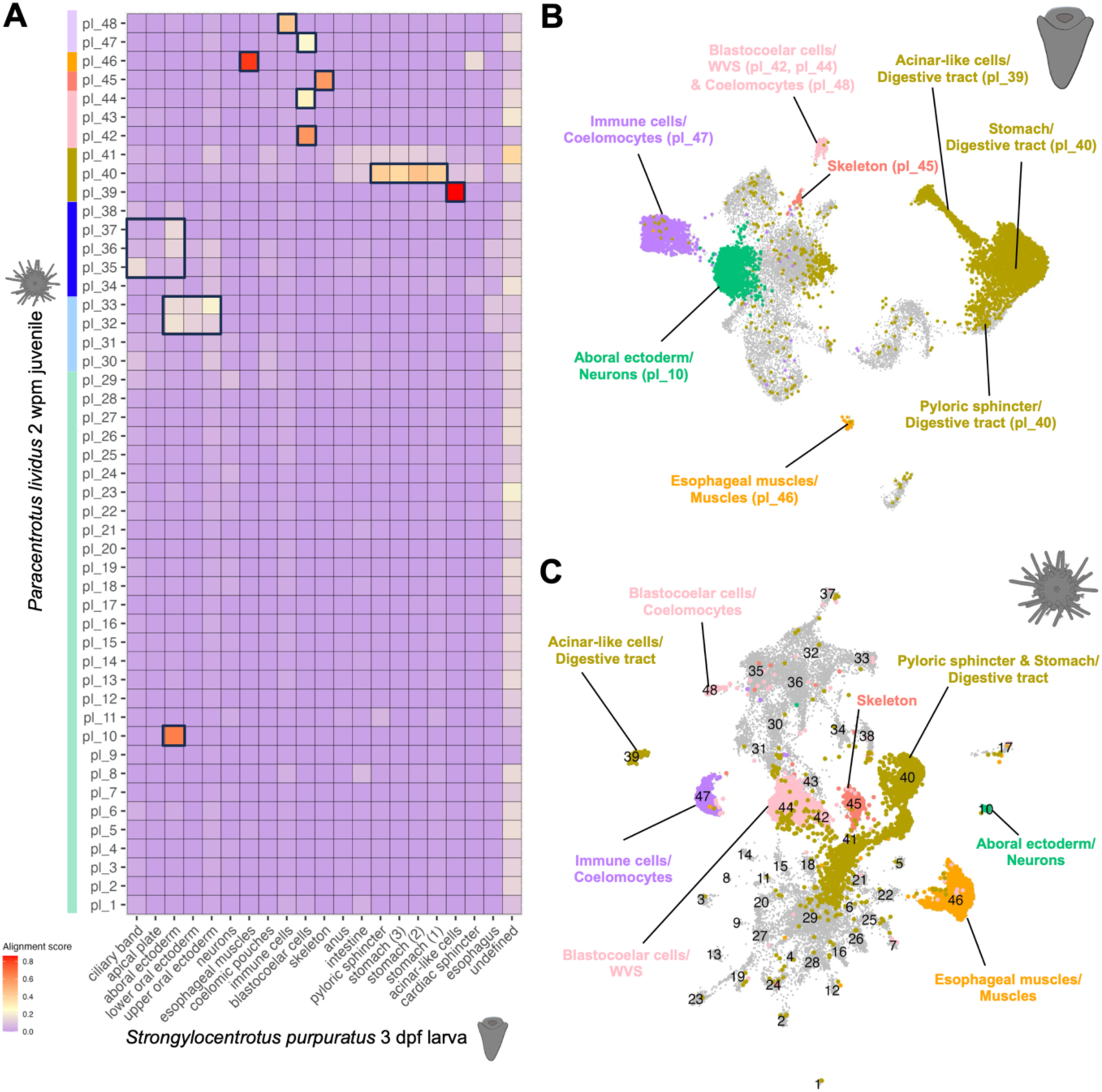
SAMap cell clusters alignment scores between *P. lividus* juvenile and *S. purpuratus* 3 dpf larva. (**A**) Mapping of the cell clusters between the snRNA-seq *P. lividus* juvenile dataset and the scRNA-seq *S. purpuratus* larva dataset. Black boxes indicate high alignment scores. Color-code for the *P. lividus* juvenile clusters is the same as in Fig. 1E. (**B**) *S. purpuratus* 3 dpf UMAP atlas projecting the high-scored *P. lividus* juvenile cell types. (**C**) *P. lividus* 2 wpm UMAP atlas highlighting the high-scored *S. purpuratus* larval cell types.

### Characterization of the *P. lividus* juvenile nervous system

Our results showing that the early larval and juvenile nervous systems are highly diverse in cell type content, prompted us to further characterize the juvenile neuronal clusters. As mentioned previously, the juvenile nervous system is composed of millions of interconnected neurons and ganglia (Fig. 4A) (37,49), and this diversity is reflected by the high number of clusters found to correspond to neurons in our snRNA-seq atlas (Fig. 4B). To characterize the identity of those neurons, we interrogated our data for the presence of neural functional gene families, including distinct neuromodulators (neurotransmitters and neuropeptides) as well as opsin genes (Fig. 4C).

**Fig. 4.**
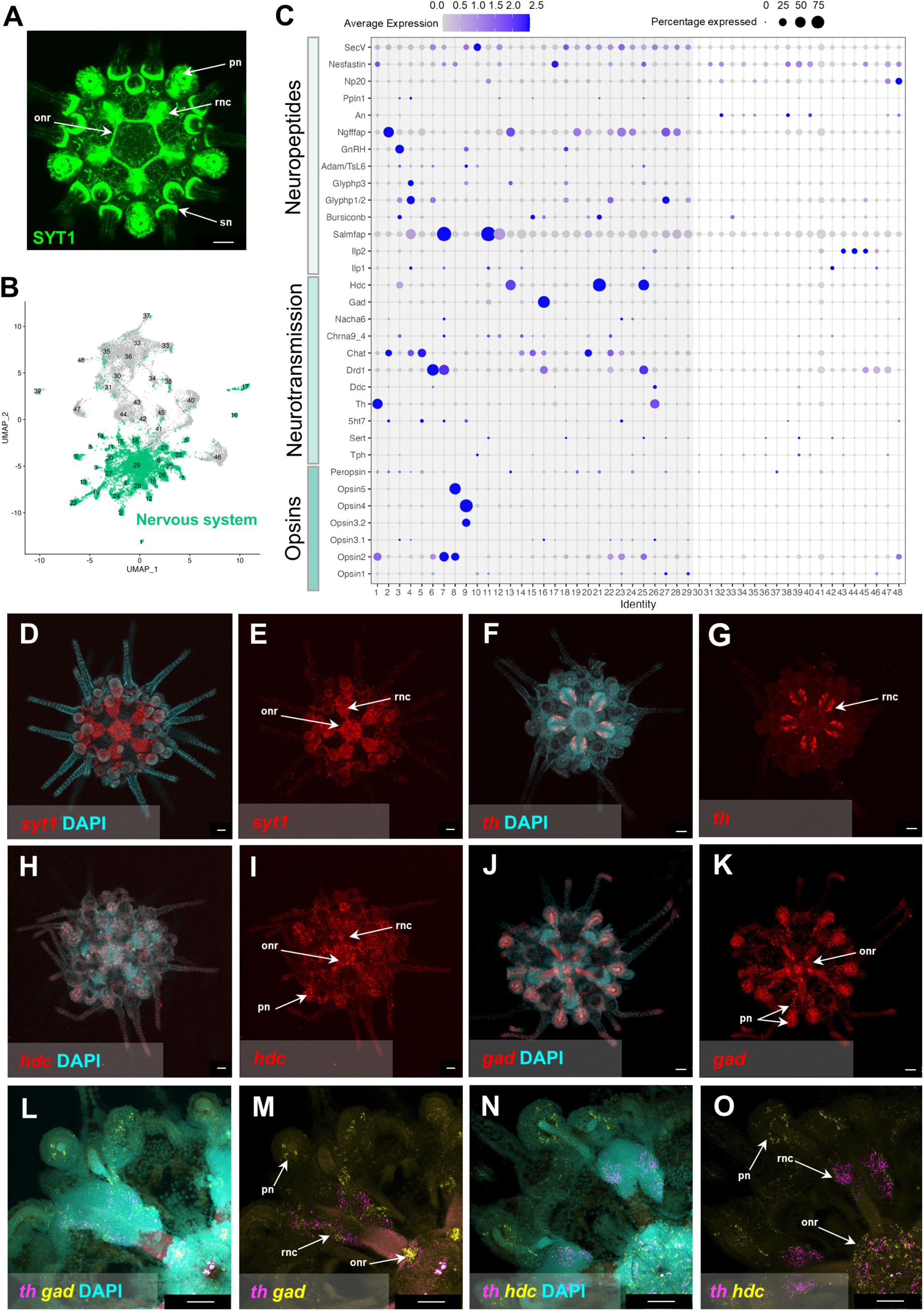
Molecular characterization of the *P. lividus 2 wpm* juvenile nervous system. (**A**) IHC for SynaptotagminB (Syt1), labeling the entire juvenile nervous system. (**B**) UMAP highlighting the neuronal clusters in the *P. lividus* juvenile snRNA-seq data. (**C**) Dotplot showing the average expression in the *P. lividus* snRNA-seq atlas of known echinoderm neuropeptides, neurotransmitters and opsin genes. (**D**) HCR for *syt1* (**D, E**), *th* (**F,G**), *hdc* (**H, I**), and *gad* (**J, K**) with (**C, F, H, J**) or without (**E, G, I, K**) nuclei staining (DAPI). (**L, M**) Double HCRs for *th* (magenta) and *gad* (yellow) and with (**L**) or without (**M**) nuclei staining (DAPI). (**N, O**) Double HCRs for *th* (magenta) and *hdc* (yellow) and with (**N**) or without (**O**) nuclei staining (DAPI). (**A**, **D-K**) Juveniles are whole-mount and in oral view. (**A, D-O**). onr, oral nerve ring; pn, podia neurons; rnc, radial nerve cord; sn, spine neurons. Scale bar is 50 μm.

Overall, we found that twenty-one out of the twenty-nine neuronal clusters contain transcripts for at least one of the five major neurotransmitter enzyme families investigated (Fig. 4C). These genes correspond to tryptophan hydroxylase (*tph*), tyrosine hydroxylase (*th*), choline acetyltransferase (*chat*), glutamate decarboxylase (*gad*), and histidine decarboxylase (*hdc*), which are respectively involved in the biosynthesis of serotonin, dopamine, acetylcholine, GABA and histamine. Interestingly, all of these twenty-one clusters also expressed at least one opsin gene, indicating a potential photosensory role. Even more striking is that except for two neuronal clusters (12 and 17) the remaining twenty-seven are all expressing opsin genes, highlighting the importance of light in the regulation of the juvenile’s physiology.

According to our snRNA-seq data, acetylcholine appears to be the most widespread neurotransmitter in the juvenile’s nervous system, with *chat* transcripts detected in twelve out of the twenty-nine neuronal clusters (clusters 2, 4, 5, 14, 15, 16, 18, 20, 22, 23, 28 and 29) (Fig. 4C). Among these clusters, the neuropeptidergic signature varies, suggesting the presence of diverse subtypes of cholinergic neurons. In the case of histaminergic neurons, we found *hdc* to be expressed in four neuronal clusters (3, 13, 21 and 25) out of the twenty-nine. Likewise, we detected *tph*, the marker of serotonergic neurons, in four of the twenty-nine neuronal clusters (10, 15, 24, 25). Searching for GABAergic and dopaminergic neurons, we identified only two clusters, for each neuronal type, expressing respectively *gad* (11 and 16) and *th* (1 and 26), although at high levels. This analysis is providing evidence of a very little overlap between the expression domains of the different neurotransmitter-related enzymes looked at, suggesting a high degree of neuronal diversity. The only overlapping predicted expression domains found are *gad* and *chat* being co-expressed in the neuronal cluster 16 and *hdc* and *tph* being co-expressed in the cluster 25.

The neuronal identity of eight clusters (6, 7, 8, 9, 12, 17, 19 and 27) could not be further defined in terms of their neurotransmitters due to the absence of enzymes involved in the biosynthesis of serotonin, dopamine, acetylcholine, GABA or histamine. For those eight clusters, we analyzed the neuropeptidergic and photoreceptor molecular signatures (Fig. 4C). For example, we found transcripts encoding for *glyph1/2, adam/tsL6, secv, opsin1* and *opsin3.1* in cluster 6; *5ht7, drd1, chrna9_4, nacha6, ilp1, salmfap, nesf, seV* and *opsin2* in cluster 7; *nesf, opsin2* and *opsin5* in cluster 8; *5ht7, glyph3, adam/tsL6, gnRH, secV, opsin3.2* and *opsin4* in cluster 9; *chrna9_4, ilp1, salmfap, ppln1* and *nesf* in cluster 12; *nesf* in cluster 17; *5ht7, gnRH, ngfffap, secv* and *peropsin* in cluster 19; *nacha6*, *ilp1*, *salmfap*, *glyph1/2*, *glyph3*, *ngfffap, secv, opsin1* and *peropsin* in cluster 27.

Spatial expression analysis, using HCR for *syt1*, an echinoderm pan-neuronal marker, and several other neuromodulators, corroborated the predictions from our snRNA-seq data. While *syt1* labeled the ONR, RNC and podia neurons (Fig. 4D, E), *th* labeled exclusively the RNC (Fig. 4F-G), and transcripts for *hdc* and *gad* were detected both in the RNC and podia neurons (Fig. 4H-K). To test whether there was any co-localization of *th* and *gad* as well as *th* and *hdc* in the RNC, we performed double HCRs for these genes. Both experiments revealed no co-expression, but complementary expression patterns in RNC neurons, thus validating the single nuclei predictions (Fig. 4L-O).

Overall, our findings provide novel insight into the neuronal types present in the post-metamorphic *P. lividus* juvenile, highlighting a high degree of nervous system diversity and showing the somewhat surprising presence of opsin expression in most of the juvenile nervous system component cell types.

### PRC signatures in the *Paracentrotus lividus* juvenile nervous system

Aiming to identify the molecular signature of a photoreceptor cell type candidate responsible for sea urchin vision out of the numerous opsin positive *P. lividus* neurons, we set out to identify the putative PRC cell types. To do so, we computationally subsetted the fraction of the neuronal cell type families that contain transcripts of at least one opsin gene, and performed further clustering analysis. This resulted in the identification of fifteen neuronal PRC clusters (Fig. 5A).

**Fig. 5.**
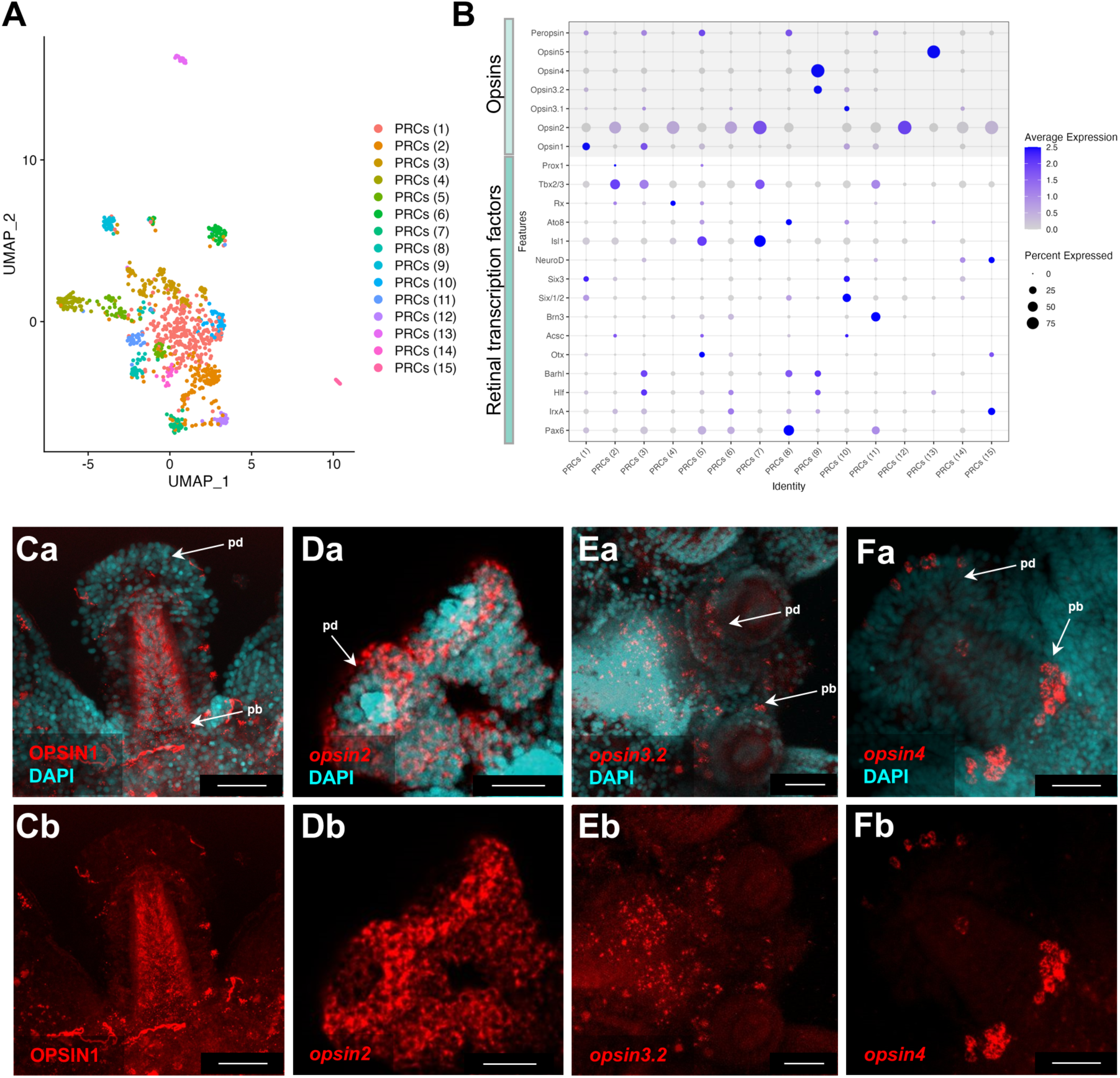
*P. lividus* 2 wpm juvenile neuronal photoreceptor repertoire. (**A**) UMAP showing the subclustered and reanalyzed opsin-positive neuronal clusters. (**B**) Dotplot showing the average expression of the opsin genes encoded in the *P. lividus* genome and of transcription factor orthologs involved in the establishment of the PRC fate in other animals. (**Ca, Cb**) IHC using a sea urchin specific opsin1 antibody. (**Da, Db**) FISH using an antisense RNA probe against *opsin2*. Nuclei are stained with DAPI (cyan). HCR using a specific probe against *opsin3.2* (**Ea, Eb),** *opsin4* (**Fa, Fb).** Nuclei are stained with DAPI (cyan). pb, podia base; pd, podia disc. Scale bar is 25 μm.

To characterize the molecular signatures of these clusters, we subsequently used genes identified from other animal taxa as essential for the specification of the PRCs and/or as important elements of the PRC identity. Such gene markers included opsins, retinal transcription factors and phototransduction molecular cascade genes. Each of them had a distinct molecular signature as found by plotting the top 10 differentially expressed marker genes (Fig. S5) and are expressing orthologs of genes that are involved in the phototransduction molecular cascade in other taxa (Fig. S6).

To understand the opsin distribution across the generated clusters, we plotted for the average expression of the seven sea urchin opsin genes within the clusters (Fig. S7), together with transcription factors whose orthologs are either essential for the specification of the PRCs and/or important elements of the PRC identity (Fig. 5B). Our analysis showed that *opsin2* was the most widely expressed opsin, i.e., being present in seven (2, 4, 6, 7, 12, 14, 15) of the fifteen reconstructed PRC clusters, followed by *opsin1*, *opsin3.1* and *peropsin* expressed in five of the fifteen PRC clusters (1, 3, 4, 10, 11; 1, 3, 6, 10, 14 and 1, 3, 5, 8, 11, respectively). *opsin3.2* transcripts were detected in only four clusters (1, 3, 9, 10), and *opsin4* and *opsin5* were found solely in cluster 9 and 13, respectively.

Moreover, our analysis revealed that many of the PRC clusters are defined by more than one opsin gene. We report the co-expression of opsins 1, 2, 3.1, 3.2 and peropsin in PRCs(1); opsins 1, 3.1, 3.2 and peropsin in PRCs(3); opsins 1 and 2 in PRCs(4); opsins 2 and 3.1 in PRCs(6); opsins 3.2 and 4 in PRCs(9); opsins 3.1 and 3.2 in PRCs(10); opsin1 and Peropsin in PRCs(11); and opsins2 and 3.1 in PRCs(14). Nonetheless, there were also some PRC clusters expressing a single opsin, such as the clusters 2, 7, 12 and 15 expressing only *opsin2*, cluster 13 that contains transcripts only for *opsin5*, and clusters 5 and 8 where only *peropsin* transcripts were detected. Transcription factors known to control the establishment and maintenance of the PRC identity in other animals were further detected in all PRC clusters. Except for the PRC cluster 12, all PRC clusters contained transcripts for more than one of these transcription factors, suggesting that PRC specification and differentiation genes are reutilized during animal evolution and that different PRC types use different transcription factor toolkits. For instance, we found *pax6*, *hlf, otx*, *six1/2*, *six3* and *isl1* co-expressed in PRCs(1); *prox1*, *tbx2/3*, *rx*, *six3*, *acsc* and *irxA* in PRCs(2); *tbx2/3, neuroD*, *barh1*, *hlf*, *irxA* and *pax6* in PRCs(3); *rx* and *isl1* in PRCs(4); *prox1*, *rx*, *ato8*, *isl1*, *acsc*, *otx*, *hlf* and *pax6* in PRCs(5); *brn3*, *otx*, *hlf*, *irxA* and *pax6* in PRCs(6); *isl1* and *tbx2/3* in PRCs(7); *ato8*, *six1/2*, *barh1*, *irxA* and *pax6* in PRCs(8); *irxA*, *hlf* and *barh1* in PRCs(9); *acsc*, *six1/2*, *six3* and *ato8* in PRCs(10); *pax6*, *brn3*, *neuroD*, *rx* and *tbx2/3* in PRCs(11); *hlf* and *ato8* in PRCs(13); *six1/2*, *six3* and *neuroD* in PRCs(14) and *six3*, *otx* and *irxA* in PRCs(15).

To check which opsin-expressing cell type cluster might correspond to a potential visual PRC candidate, we analyzed spatially the expression domains of some of the opsin genes expressed by the *P. lividus* 2 wpm juveniles using IHC for opsin1, FISH for *opsin2* and HCR for *opsin3.2* and *opsin4*. We found *opsin1* expression in the podia (Fig. 5Ca,Cb) as well as in various ectodermal tissues, consistent with previous observations (37). By comparison, *opsin2*, *opsin3.2* and *opsin4* were confined to the podia domain (Fig. 5Da-Fb). *opsin3.2* and *opsin4* transcripts were found in overlapping domains at the base and in the disc of the juvenile podia respectively, with additional *opsin3.2* positive cells being present in the surrounding epidermis (Fig. 5Da-Fb). Those expression patterns corroborate the snRNA-seq predictions showing *opsin3.2* being co-expressed with *opsin4* in the PRC cluster 9 (Fig. 5B).

Reconstructing the PRC map of the sea urchin juvenile nervous system, along with the spatial expression patterns of opsin genes, enabled us to refine our search for cell type candidates that could mediate the animal’s visual behaviors. While PRCs occur widely throughout large portions of the animal’s epidermis and within other tissues, including those expressing *opsin1, opsin2* and *opsin3.2*, these cells likely mediate a variety of physiological functions requiring photoreception (53). However, sea urchin vision has been hypothesized to depend on specific PRC “units”, as suggested by the first neuro-mathematical model of sea urchin vision (54).

From our analysis, only two PRC clusters matched these criteria: the PRC cluster (13) positive only for *opsin5* and the PRC cluster (9 exclusively positive for *opsin4* (Fig. 5B). *opsin5* codes for an echinoderm-specific opsin type, named Echinopsin, which has currently no known function in sea urchins and is phylogenetically distinct from any of the opsin types typically involved in metazoan vision (55). In contrast, melanopsin-expressing PRCs have previously been hypothesized to facilitate image-forming vision in sea urchins (25,54). Our analysis revealed that a single PRC subcluster expresses melanopsin (opsin4) in the *P. lividus* juvenile (Fig. 5B, Fig. S7). *Opsin4* mRNA and protein were detected exclusively in two morphologically distinct cell aggregations: one at the base (Fig. 6A) and another within the disc of the juvenile’s podia (data not shown). In both aggregations, we found co-expression of *opsin4* and *opsin3.2*. To characterize the identity of the opsin4/opsin3.2 double-positive cell type, we plotted the average expression of the top 50 differentially expressed marker genes of PRC(9) (Fig. 6B). Among these genes, we detected transcripts for *opsin4, opsin3.2, beta-arrestin, cryptochrome1 (vcry)*, and *nitric oxide synthase*, all of which are known components of phototransduction and UV avoidance pathways.

**Fig. 6.**
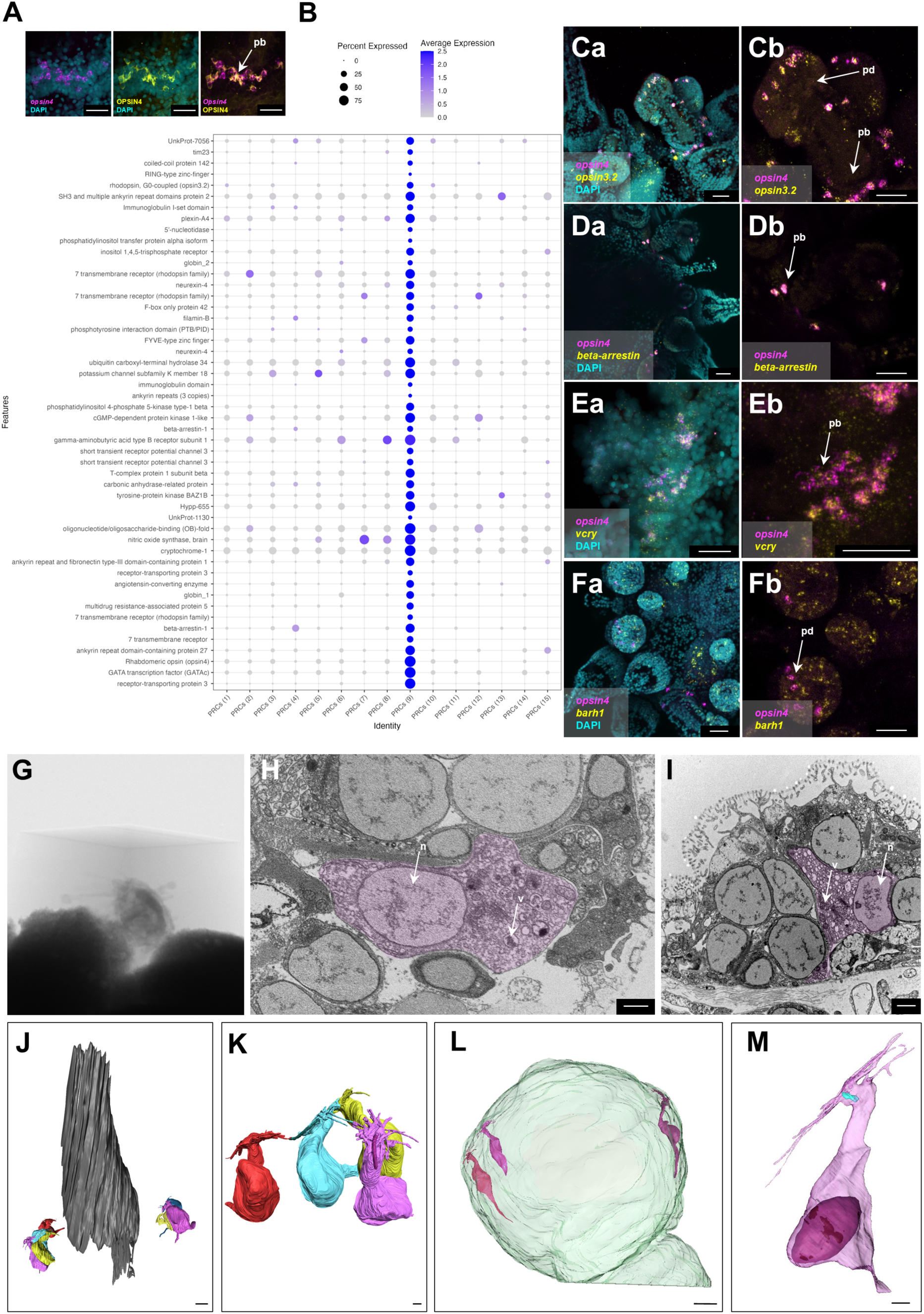
Molecular and morphological reconstruction of the *P. lividus* 2 wpm juvenile opsin4 (melanopsin) cell type. (**A**) HCR for *opsin4* (magenta) paired with IHC using a sea urchin specific opsin4 antibody (yellow). Nuclei are stained with DAPI (cyan) (**B**) Dotplot showing the average expression of the top 50 marker genes of the PRC(9) cell cluster. Double HCR using a specific probe against *opsin4* paired with *opsin3.2* (**Ca, Cb),** *arrestin* (**Da, Db),** *vcry* (**Ea, Eb)** and *barh1* (**Fa, Fb).** Nuclei are stained with DAPI (cyan). (**G)** Overview of one of the embedded 2 wpm *P. lividus* specimen before being processed for SBF-SEM. (**H)** SBF-SEM isolated section showing the opsin4-positive PRCs at the base of the podia. (**I)** TEM isolated slice showing the opsin4-positive PRCs at the podia disc. (**J)** 3D reconstruction of the opsin4-positive PRCs at the base of the podium. podium 3D segmented membranes depicted in gray. (**K)** Close-up of the 3D reconstructed basal PRCs. (**L)** 3D reconstruction of the podium disc PRCs. Disc 3D segmented membranes are in pale green. (**M)** Close-up of a 3D reconstructed disc PRC. Ciliary rootlet is labelled in pale blue. n, nucleus; pb, podia base; pd, podia disc; v, vesicle. Scale bars in panels A, Ca-Fb are 25 μm, in J 4 μm while in H, I and K-M 1 μm.

To validate the spatial co-expression of genes predicted by our snRNA-seq analysis, we performed double HCR on whole-mount *P. lividus* 2 wpm juveniles. We confirmed co-expression with *opsin4* for *opsin3.2* (Fig. 6Ca, Cb), *beta-arrestin* (Fig. 6Da, Db), *vcry* (Fig. 6Ea, Eb), and *barh1* (Fig. 6Fa, Fb). Interestingly, in all of our in situ validations, co-expression was detected in both opsin4 expression domains—the base and in the disc of the juvenile’s podia. Given that our snRNA-seq analysis consistently identified only a single cluster containing opsin4 transcripts (PRC cluster 9) (Fig. 5B), this suggests that the opsin4-positive cells at the base and in the disc of the podia belong to the same cell type family.

To further investigate whether juvenile *P. lividus* possesses only one melanopsin-positive cell type or whether there is a convergence of molecular PRC signatures, we examined the morphological characteristics of these two cell populations using serial block-face electron microscopy (SBF-SEM) and high-resolution transmission electron microscopy (TEM). Serial sectioning of resin-embedded *P. lividus* juveniles (Fig. 6G) revealed enlarged microvilli-bearing PRCs in both the basal podia region and the podia disc. Densely packed, large membrane vesicles inside the PRC cytoplasm, which have been shown to carry melanopsin in *Strongylocentrotus purpuratus* (25), were clearly distinguishable in both the basal podia (SBEM, Fig. 6H) and the podia disc regions (TEM, Fig. 6I).

In *P. lividus* 2 wpm juveniles, we further identified at the base of the podium 2 bilaterally symmetrical PRC aggregations, each containing three to four PRCs (Fig. 6L, N), consistent with previous immunostainings against sea urchin melanopsin in the developing rudiment of *P. lividus* (37), we here identified in 2 wpm juveniles two bilaterally symmetrical PRC aggregations flanking the podium base, each containing three to four PRCs (Fig. 6H, J). A 3D reconstruction of their morphology revealed flask-shaped cells extending a thin dendrite and numerous long microvilli projecting from the apical cell surface (Fig. 6K). Some PRCs bore an unmodified cilium (Fig. 6K), meeting the structural requirement for a Go-Opsin (*opsin3.2*) potentially functioning in those PRCs. In cells lacking a cilium, a ciliary rootlet was observed, suggesting that the cilium might have been lost during chemical fixation. At this developmental stage of *P. lividus,* we observed only two to three PRCs in each podium disc (Fig. 6L, M) forming epidermal patches with other ciliated cells with presumed receptor function. Except for the differing spatial PRC distribution (single PRCs in the podium disk versus PRC aggregations at the podium base), the cells feature an identical morphology and ultrastructure.

## Discussion

### Signature, conservation, and divergence of pre- and post-metamorphic cell types

Analyzing the cell type atlas of post-metamorphic juvenile stages of the sea urchin *P. lividus*, enabled us to assess questions such as: What are the cell types constituting a post-metamorphic sea urchin juvenile? How similar are those cell types to the larval ones? What components constitute the juvenile nervous and photoreceptive systems? Previous studies using molecular markers have already reported on the distribution of juvenile muscle, skeletal, and neuronal cell types in post-metamorphic developmental stages (37,45,49,50). Our analysis confirmed these findings at the single-cell level and provided additional knowledge on these cell types as well as disclosed the molecular signatures for five additional major cell types. Altogether, our snRNA-seq survey highlighted the molecular signatures of eight major *P. lividus* cell type groups: muscles, neurons, epidermis, podia epidermis, skeleton, coelomocytes, water vascular system, and digestive tract, all of which are distributed across forty-eight distinct cell type clusters.

At the regulatory signature level, the only available data regarding post-metamorphic juveniles pertain to the skeleton. Previous studies have shown that a few of the genes constituting the embryonic and larval skeletogenic GRN (*vegfr, alx1, sm37, sm50, mps130*) are expressed in the juvenile skeletal cells, suggesting a partially conserved skeletogenic regulatory program between embryos, larvae and juveniles (45,50). Our findings here corroborate this statement. Indeed, our study reveals that, with the exception of four genes, almost the entire embryonic/larval reconstructed skeletogenic GRN, comprised of twenty-six genes, including the regulatory genes and the differentiation gene batteries, operates in post-metamorphic skeletal cells, exemplifying how a cell type can be formed *de novo* using conserved genomic and regulatory information.

Overall, our analysis demonstrates that many juvenile cell types show molecular similarity to their larval counterparts. For instance, juvenile muscle cells exhibit high molecular similarity to larval ones, utilizing most of the regulators involved in the embryonic myogenesis program. Furthermore, we identified distinct clusters of juvenile coelomocytes that align with either larval immune cells, a cell type family that contains globular and pigment cells (9), or larval blastocoelar cells, all of which are components of the sea urchin pre-metamorphic immune system (56). Notably, the alignment of larval blastocoelar cells with juvenile WVS clusters further suggests that the larval blastocoelar cell network may function akin to a vascular system.

Interestingly, of the three juvenile digestive tract clusters identified in our analysis, only one corresponds to the larval mid/posterior gut, one to the larval acinar-like cells and one appears to be specific to the juvenile stage. Notably, we found no alignment between the larval clusters corresponding to the anus, intestine and foregut with any of the juvenile digestive tract clusters, suggesting that while similar transcription factors are expressed in those cells the cell type outcome is different. This is not so surprising given that these larval tissues are known to be reabsorbed during the metamorphic process (57). Another explanation for the lack of similarity between those cell types could be the different feeding strategies of sea urchin larvae and juveniles feeding on microorganisms and algae, respectively. Such differences must dictate the presence of diversified gut cell types involved in proper food digestion and nutrients uptake. For acinar-like cells, our previous research demonstrated their molecular and morphological homology to the exocrine acinar cells of mammals (43). Their presence in juveniles represents the first evidence of a pancreatic-like cell type in post-metamorphic echinoderms. While our findings indicate here that most juvenile cell types are formed *de novo*, the larval stomach and acinar-like cells may be retained and used as a scaffold for completing juvenile gut organogenesis, as suggested by previous morphological studies (57). Thus, the digestive tract represents a particular example of cell types mostly inherited from the larva, rather than newly formed in juveniles.

### The echinoderm germline

Interestingly, in our snRNA-seq atlas no cluster corresponding to the germline could be identified. It has previously been postulated that sea urchins possess an inherited mode of germ line specification, based on the expression of gene markers commonly found in germ cells and somatic stem cells in various animals (58–60). However, the functional role of these markers remains unclear. It is important to note that this cell population corresponds to small micromere descendants, which during embryogenesis populate the foregut region and, during larval development, are detected within the coelomic pouches (9,61).

Both our gene candidate approach and SAMap analysis failed to identify a juvenile cluster matching the coelomic pouches cluster of the larva, where cells with germ/stem molecular signatures have been reported (9,61). Moreover, germ/stem cell molecular signatures were found scattered in a broad spectrum of clusters corresponding to somatic cell types. Several scenarios could explain this observation. A first possibility is that the germline cells may be present in juveniles but simply few in number or embedded within deeper tissue layers that our dissociation protocol failed to extract. In disagreement with this hypothesis is, however, the fact that we retrieved in our snRNA-seq atlas rare cell types, such as opsin4-positive cells, as well as digestive tract-related cell types, which are located in the deepest layers of the juvenile body cavity. Another possibility is that the embryonic and larval germ/stem cell populations represent stem cells rather than germ cells. This hypothesis is supported by their location in the coelomic pouches, a domain where the juvenile rudiment forms. These cells could thereby contribute to juvenile formation rather than directly to the germline, and consequently, the germline could be specified later, during juvenile or adult development.

### Evidence of an “all-brain” echinoderm state

The most abundant and diverse group of cell types in terms of cluster representation, as revealed by our analysis, corresponds to neurons. While we identified the expression of most embryonic and larval neurogenic patterning genes in the juvenile nervous system clusters, we found no direct similarity between larval and juvenile neurons. This finding suggests that although the same genetic toolkit is used to generate neurons, the outcomes of the neurogenic program differ significantly between the two analyzed life stages. The only exception is a single juvenile neuronal cluster that exhibited similarity to an ectodermal larval cluster. This result reflects a convergent molecular signature of patterning genes. Consistent with this, it has been previously shown that the aboral ectoderm cells of sea urchin larvae express several genes also involved in the specification of the larval apical plate (9) and that neurogenesis occurs in the aboral size of the animal plate (62). Alternatively, it is also possible that as for the digestive tract, some larval neurons present in the aboral ectoderm have been inherited by the juveniles, during metamorphosis.

Traditionally, the echinoderm nervous system has been characterized as “simple,” due to the absence of a centralized brain. It was previously postulated that the expression of key CNS orthologs in the adult sea urchin pentaradial nervous system is indicative of a five-fold interconnected CNS duplication (63). The large number of neuronal clusters we identified, comprising more than half of the juvenile cell atlas clusters, and the high diversity of their molecular signatures (including serotonergic, histaminergic, dopaminergic, GABAergic, cholinergic, and various neurosecretory neurons) indeed suggest a far more sophisticated nervous system than a mere network of interconnected neurons and ganglia. Adding to the nervous system complexity, we found a widespread distribution of neuronal clusters with an ANE molecular signature, that in marine invertebrate larvae is required for the formation of neurons with an analogous role to the vertebrate CNS, thus suggesting an anterior or brain-like organization of the echinoderm nervous system. In line with this hypothesis, we show that the ANE GRN genes *nkx2*.1 and *frz5/8* are expressed in both ONR and RNC that comprise the largest part of the sea urchin adult nervous system. Moreover, a recent study has demonstrated that in chordates an orthologous subset of the sea urchin ANE genes, including *nkx2.1* and *frz5/8* constitute the anterior GRN that controls the forebrain identity (52). Further, corroborating our hypothesis, a recent study on the sea star *Patiria miniata* juvenile demonstrated that echinoderms are predominantly “head-like” animals in terms of ectodermal patterning (40), while similarly *nkx2.1* and *frz5/8* were found expressed in the same nervous system domains as the ones we report in this study. Additionally, the anterior neuroectoderm patterning genes *sfrp1/5, frz5/8, six3* and *nkx2.1* were reported to be expressed in the ONR and RNC in the post-metamorphic brittle star juvenile (64). Taken together these data suggest that the brain-like organization hypothesis could apply to the entire echinoderm clade. Therefore, our data, which demonstrate the expression of several vertebrate CNS homologs in tissues throughout the juvenile nervous system, not only support the “head-like” hypothesis for echinoderm axial patterning, but also suggest that the nervous system of these allegedly “brainless” animals features an “all-brain’’ organization.

### Melanopsin and Go-opsin are co-expressed in a deuterostome photoreceptor

Adding to the molecular complexity of the sea urchin nervous system is the surprising finding that most neuronal cell types express at least one opsin gene. This suggests that large portions of the juvenile and adult sea urchin nervous system are under light-dependent control. Fifteen clearly distinguishable photoreceptor putative cell (PRC) types, each expressing a specific combination of evolutionarily conserved retinal transcription factors, highlight the remarkable complexity of the sea urchin photoreceptor system.

Despite the numerous opsin-expressing neurons and in contrast to the extensive gene duplications observed in melanopsin-expressing PRCs of brittle stars (65,66) and crinoids (67), our snRNA-seq data revealed the expression of only seven specific opsin genes in the juvenile *P. lividus*, with no evidence of opsin gene duplication. Apart from opsin2-expressing PRCs, which have recently been shown to modulate larval swimming behavior in the sea urchin *Hemicentrotus pulcherrimus* (68,69) the specific functions of most echinoderm opsins remain poorly understood.

PRCs involved in animal vision predominantly express either a ciliary opsin (localized inside often-modified PRC cilia) or a melanopsin/rhabdomeric opsin (localized inside often-modified PRC microvilli) (70). Our spatial expression analysis of the sea urchin ciliary opsin in juvenile *P. lividus* corroborates previous findings in other sea urchins where ciliary opsin is expressed across the entire juvenile body (37,71), rendering it an unlikely candidate for sea urchin vision. In contrast, *P. lividus* melanopsin (opsin4)-expressing PRCs exhibit a similar localization and ultrastructure to those described in *S. purpuratus* (25). PRCs at the podial base and disc were morphologically indistinguishable from one another; a finding in accordance with our molecular analysis showing only one single melanopsin-positive PRC subcluster. We thus propose only one melanopsin-expressing PRC type in *P. lividus* and potentially in other sea urchins, which have shown comparable spatial melanopsin expression patterns (72).

Interestingly, *P. lividus* melanopsin-expressing PRCs co-express a Go-opsin (opsin3.2), a type of opsin previously reported in only a small number of marine invertebrates (55,73–75). In sea urchins, the Go-opsin has so far only been documented in larval cells. Valero-Gracia and colleagues first identified it in two cells flanking the larval apical organ (76). Valencia and colleagues thereafter reported some transcription factors typically associated with ciliary PRCs, while other typical ones were absent (77). Cocurullo and colleagues also identified Go-opsin within modified cilia of neurons (28). Finally, Yaguchi & Yaguchi demonstrated a role for Go-opsin3.2 in light-dependent pyloric opening in sea urchin larvae (78).

Our discovery of a PRC co-expressing melanopsin and Go-opsin represents a novelty within deuterostomes. Go-opsin-expressing PRCs seem to be absent in vertebrates and have so far only been found in the chordate *Amphioxus* (79,80), though no spatial expression or functional data is available to date in these animals. Phylogenetic analyses place sea urchin Go-opsins within the same clade as those found in the marine polychaete worm *Platynereis dumerilii* (53), for which Gühmann and colleagues reported co-expression of a Go-opsin with melanopsin in larval eye photoreceptors (73). Here, the Go-opsin mediates spectral tuning of the melanopsin photopigment, thereby enhancing PRC efficiency. Given the dimly lit marine environments inhabited by sea urchins such as *P. lividus*, similar spectral tuning could certainly benefit their visual capabilities.

While the presence of Go-opsin-expressing PRCs at the base of Bilateria is undisputed (53), our findings provide the first evidence of such an opsin being co-expressed with a canonical melanopsin in deuterostomes. This also underscores that although the same opsin genes, in this case *opsin3.2*, are expressed in both pre- and post-metamorphic animals, their functional roles may differ significantly.

By comparing cell type gene expression profiles between sea urchin larvae and juveniles at a single cell level, we here demonstrate the developmentally conserved and diverse building blocks of the post-metamorphic body plan. The complexity of the sea urchin nervous system, as characterized by the diversity of post-metamorphic neuronal cell type signatures and their integration of diverse PRC systems, leads us to propose that the sea urchin nervous system in its entirety comprises an “all-brain” rather than a “no-brain” state. Different PRC systems, including the proposed visual PRCs co-expressing a melanopsin and a Go-opsin feed into this complex “all-brain” network playing an indispensable role in sea urchin photobiology.

## Acknowledgements

The authors would like to thank Davide Caramiello (SZN) for animal maintenance and for providing the algae used to feed the sea urchin larvae. They would also like to thank the Service Moyen de la Mer, the Service Aquariologie and the Plateforme d’Imagerie par Microscopie of the IMEV for their support with the animals and image acquisitions. The authors are grateful to Paolo Ronchi for his assistance on the SBEM acquisitions. We would like to thank Mathilde Paris for helpful recommendations regarding the mapping of the snRNA-seq data and Michalis Averof for hosting us in his lab at IGFL, where the snRNA-seq were conducted. We are also grateful to Rossella Annunziata (SZN) for kindly providing feedback on our manuscript.

## Author contributions

Animal provision and juvenile cultures: FC, JCC, MC, OA; isolation of intact nuclei and single nucleus RNA sequencing: AA, PP; sequencing data analysis and bioinformatics: DV, PP; juvenile fixation and sample preparation for EM: FC, GB, JU-L; EM segmentation and 3D reconstructions: AZ, BZ, JC, JU-L; immunohistochemistry: JU-L, MS, PP; fluorescent in situ hybridization: PP; hybridization chain reaction (HCR): JCC, JU-L, MLR, MS, OA, AZ, TS; conceptualization and design of the project: PP, JU-L, CL, MIA; project coordination and supervision: JU-L, MIA; funding acquisition: CL, MIA; figure preparation: PP, JU-L; manuscript writing – original draft: JU-L, PP; manuscript writing – review & editing: CL; DV; FC, GB, JC, JCC, JU-L, MC, MIA, MLR, MS, OA, PP, TS. All authors revised and approved the manuscript.

## Supplementary figures

**Figure S1.**
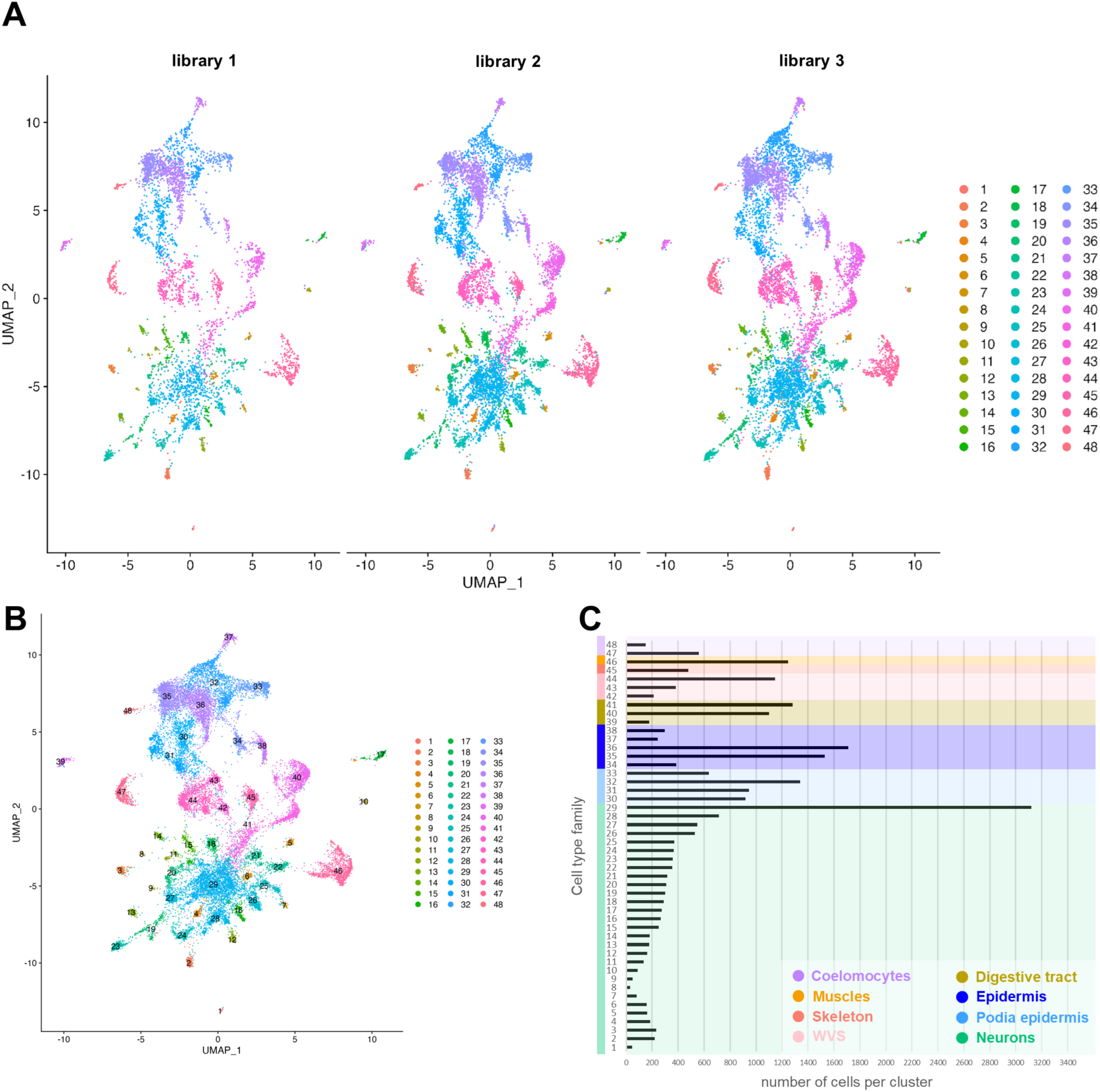
Overlap of the three snRNA-seq *P. lividus* 2 wpm juvenile libraries and cell distribution. (**A**) UMAPs of the integrated snRNA-seq data split by original identity. Cluster color code is provided on the right. (**B**) UMAP of the integrated snRNA-seq data. Cluster color code is as in panel A. (**C**) Distribution of cells per cluster.

**Figure S2.**
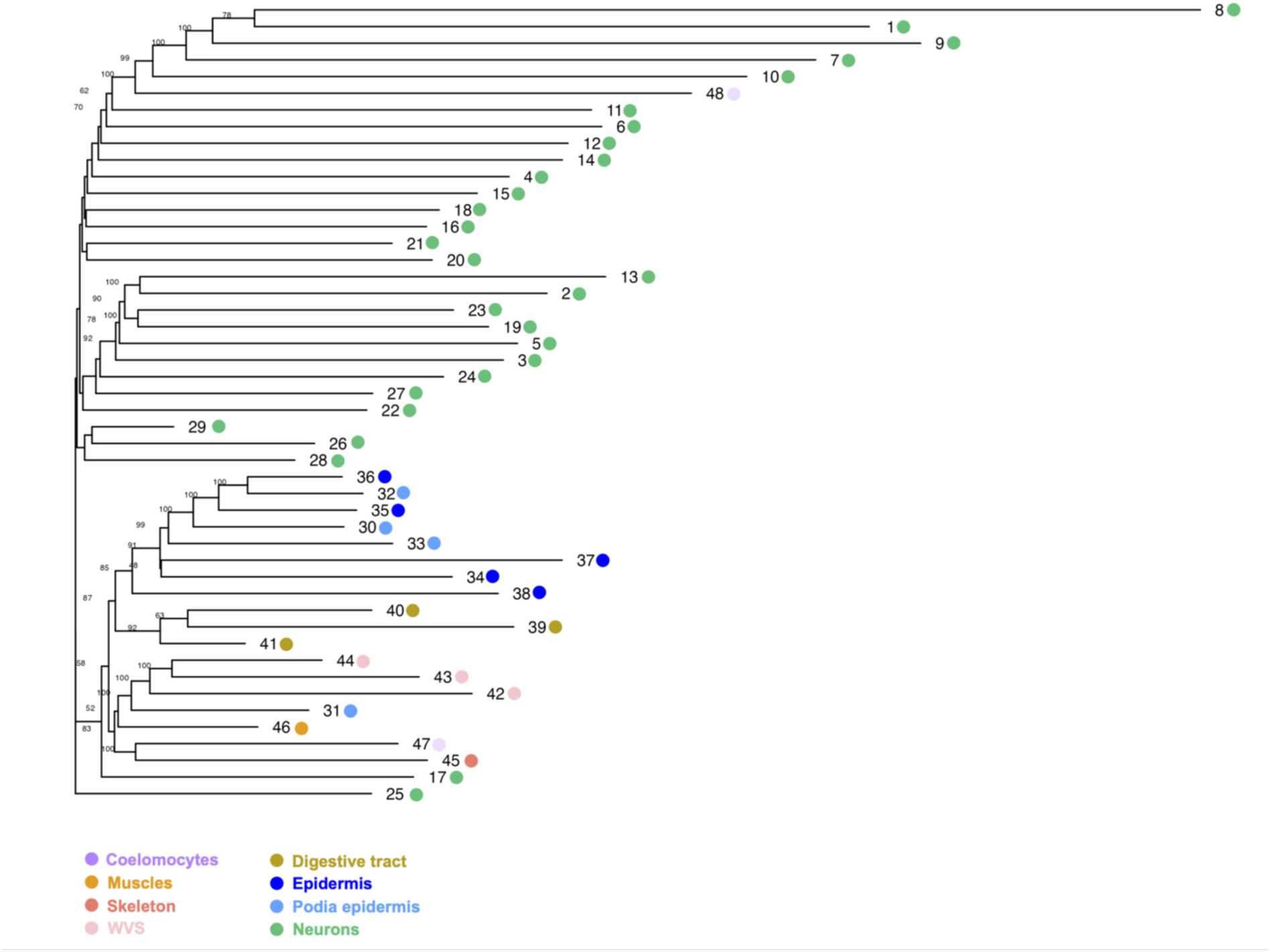
Cell type tree for the *P. lividus* 2 wpm juvenile. Cell type tree reconstructed using all expressed genes.

**Figure S3.**
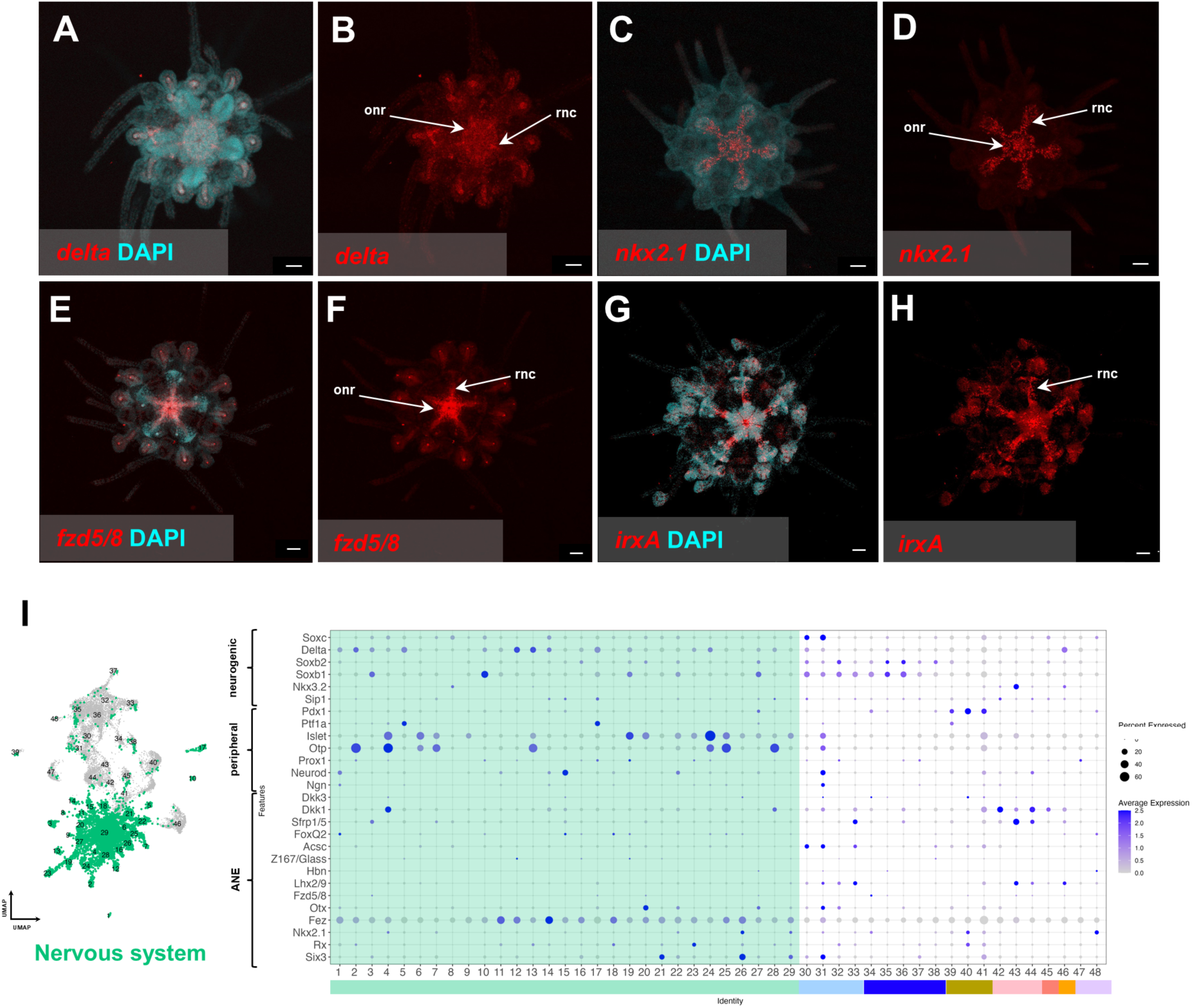
Expression patterns of neuronal signaling molecules and transcription factors in 2 wpm *P. lividus* juveniles. (**A-B**) HCR for *delta* with (**A**) and without (**B** nuclei labelling (DAPI). (**C-D**) HCR for *nkx2.1* with (**C**) and without (**D**) nuclei labelling (DAPI). (**E-F**) HCR for *fzd5/8* with (**E**) and without (**F**) nuclei labelling (DAPI). (**G-H**) HCR for *irxA* with (**G**) and without (**H**) nuclei labelling (DAPI). (**I**) Dotplot showing the average expression of the sea urchin embryonic and larval ANE, peripheral neurons and neurogenic gene markers used to score and generate panel Figure 4 panel D. (**A-H**) Juveniles are whole-mount and in oral view. onr, oral nerve ring; rnc, radial nerve cord. Scale bar is 50 μm.

**Figure S4.**
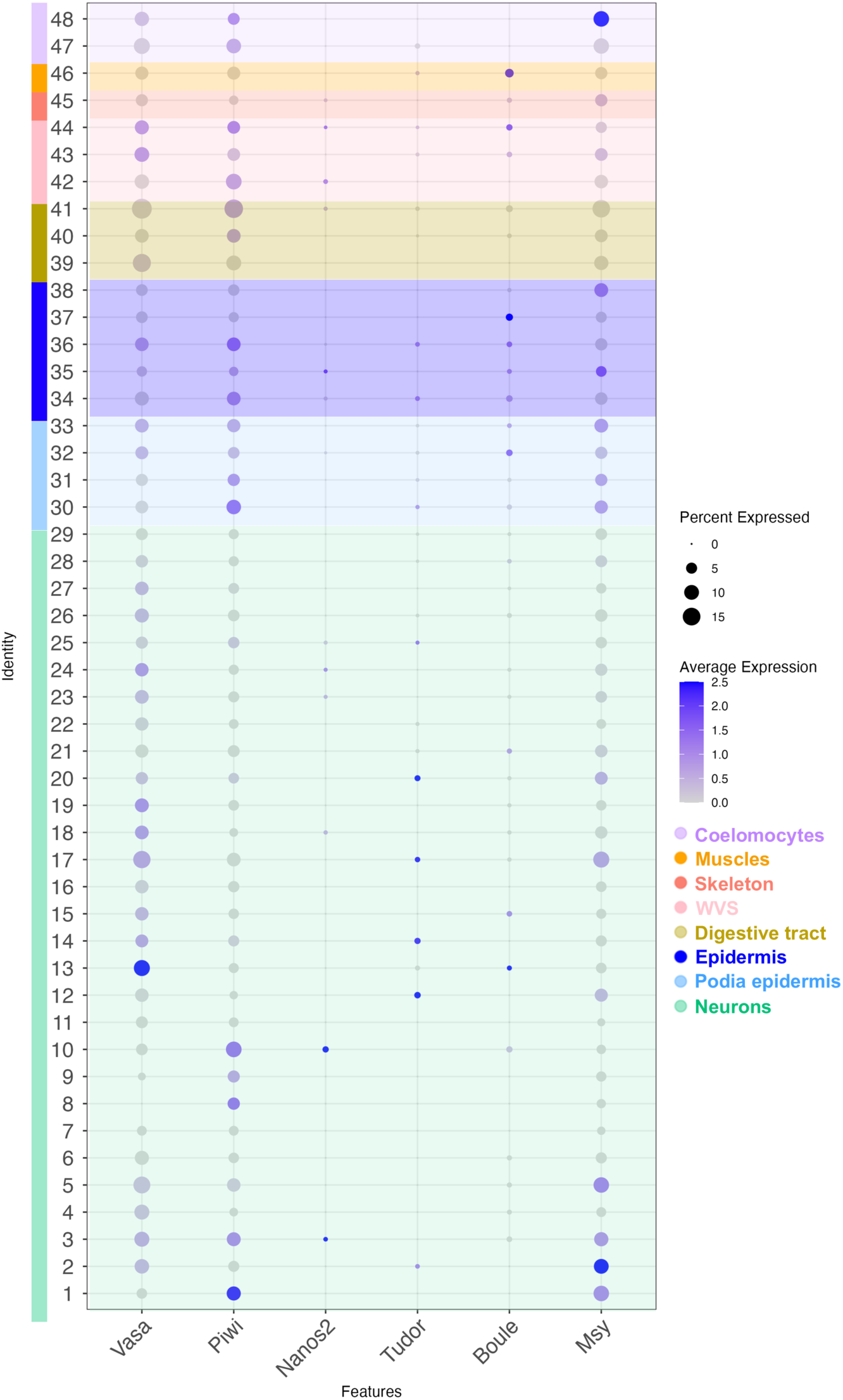
Germ/stem cell molecular markers in *P. lividus* 2 wpm juvenile. Dotplot showing the average expression of *P. lividus* orthologs of genes known to be expressed in germ/stem cells in other animals.

**Figure S5.**
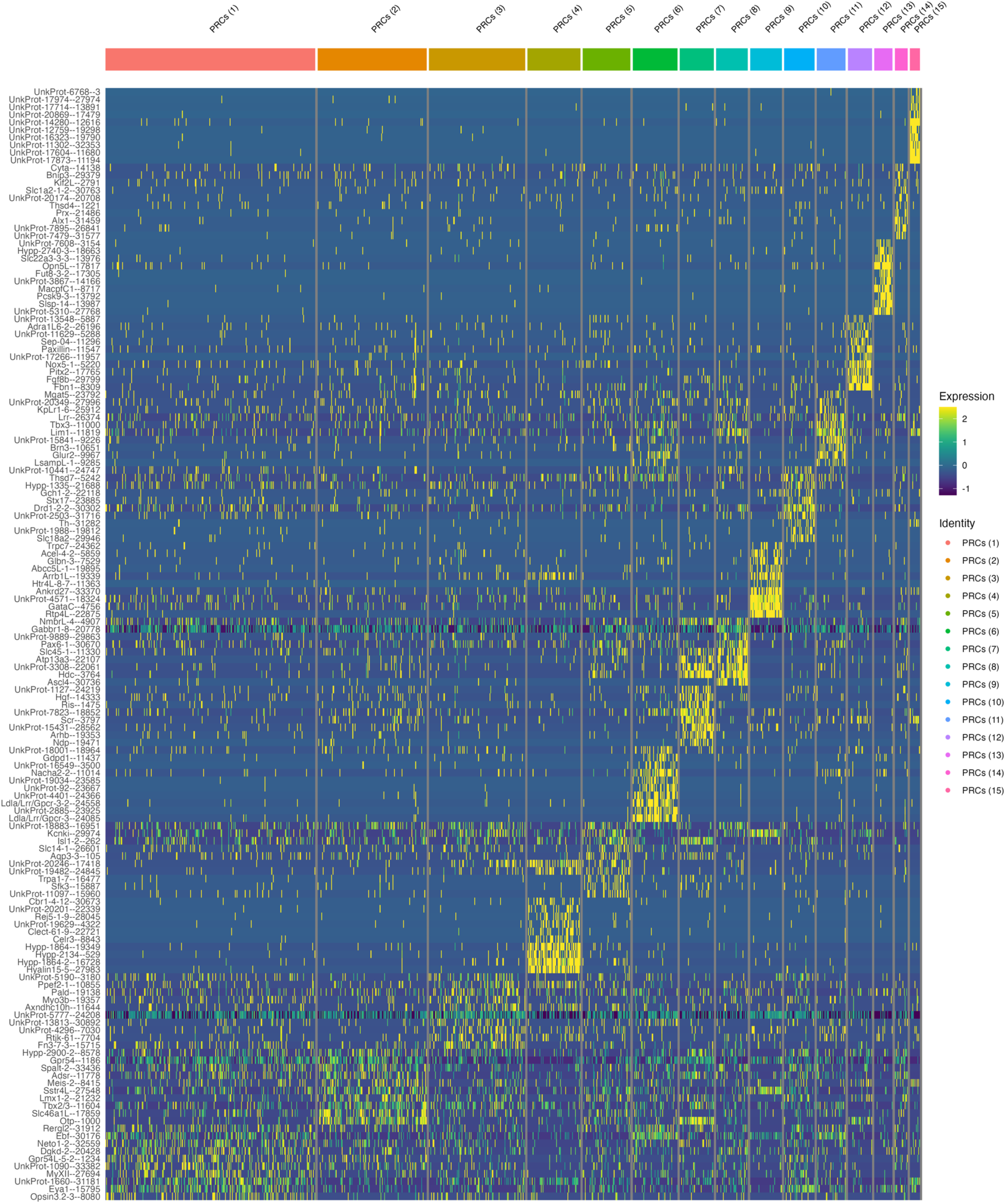
Molecular fingerprint of the *P. lividus* 2 wpm juvenile PRC clusters. Heatmap showing the top 10 marker genes of each of the PRC clusters. PRC cluster color-code is as in Figure 5.

**Figure S6.**
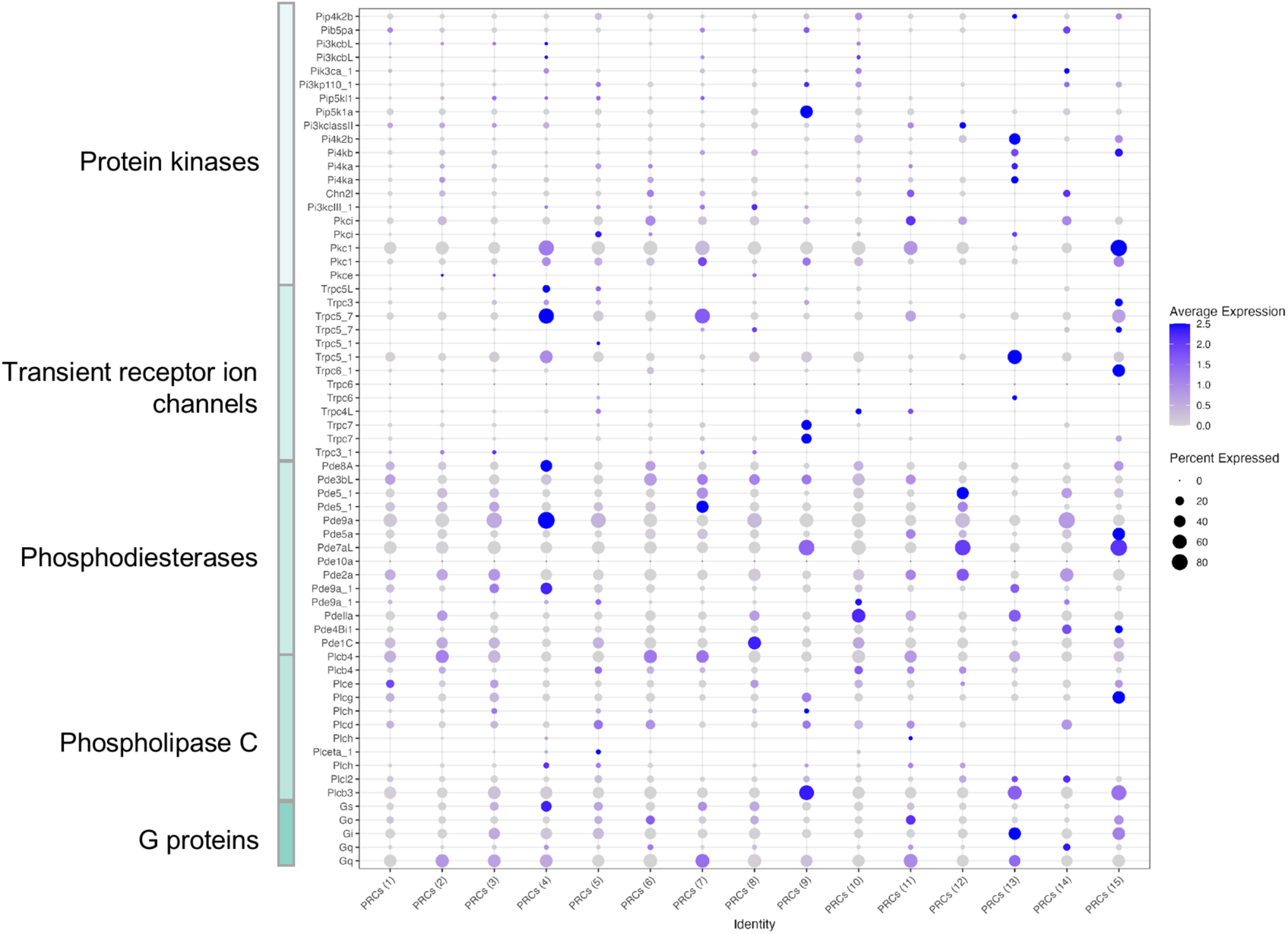
Phototransduction molecular elements in *P. lividus* 2 wpm juvenile PRC clusters. Dotplot showing the average expression of *P. lividus* orthologs of genes known to be involved in phototransduction cascades in other animals.

**Figure S7.**
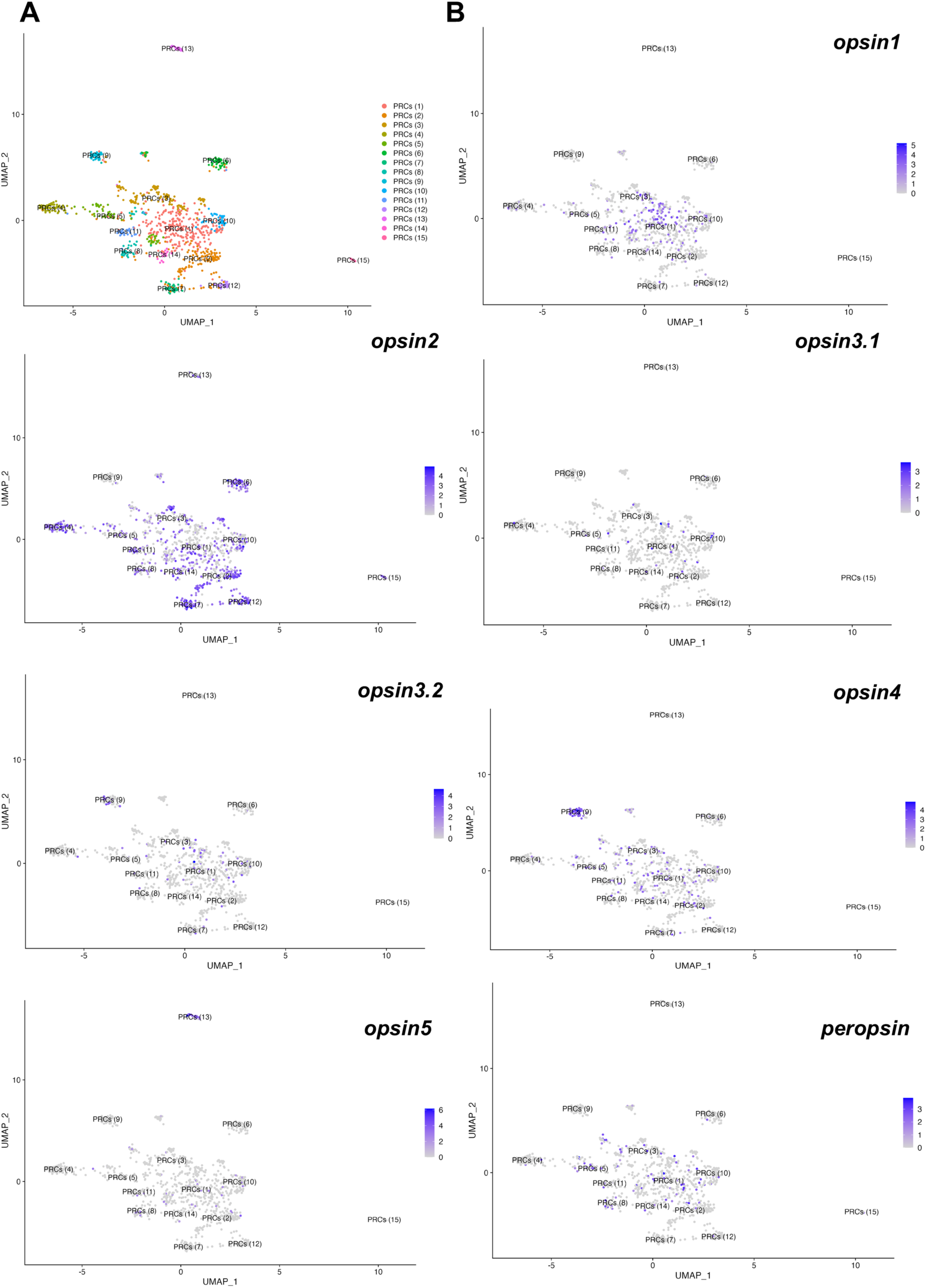
Opsin distribution across the *P. lividus* 2 wpm juvenile PRC clusters. (**A)** UMAP showing the PRC neuronal clusters. (**B)** Featureplot showing the expression of the seven *P. lividus* opsin genes within the PCR neuronal clusters.

